# Specific type 1 diabetes risk genes underpin age-at-diagnosis and indicate joint defects in immunity, beta-cell fragility and responses to viral infections in early-onset disease

**DOI:** 10.1101/577304

**Authors:** Jamie R.J. Inshaw, Antony J. Cutler, Daniel J.M. Crouch, Linda S. Wicker, John A. Todd

**Affiliations:** JDRF/Wellcome Diabetes and Inflammation Laboratory, Wellcome Centre for Human Genetics, University of Oxford, United Kingdom

**Author notes:** **Corresponding authors:** Mr Jamie Inshaw, Professor John Todd, JDRF/Wellcome Diabetes and Inflammation Laboratory, Wellcome Centre for Human Genetics, NIHR Oxford Biomedical Research Centre, Nuffield Department of Medicine, Roosevelt Drive, Oxford, OX3 7BN, 01865 287859.

## Abstract

**Objective:** Immunohistological analyses of pancreata from patients with type 1 diabetes suggest a stratification of islet pathology of both B and T lymphocyte islet inflammation common in children diagnosed at <7 years (<7 group), whereas B cells are rare in those diagnosed age ≥13 (≥13 group). Based on these observations, we sought to identify differences in genetic susceptibility between these age-at-diagnosis groups, to inform on the aetiology of the most aggressive form of type 1 diabetes that initiates in the first years of life.

**Research Design and Methods:** Using multinomial logistic regression models, we tested if known type 1 diabetes loci (17 within the HLA region and 55 non-HLA regions) had significantly stronger effect sizes in the <7 group compared to the ≥13 group, using genotype data from 27,075 individuals (18,488 controls, 3,109 cases diagnosed at <7, 3,754 at 7-13 and 1,724 at ≥13).

**Results:** Six HLA haplotypes/classical alleles and seven non-HLA regions, one of which functions specifically in beta cells (*GLIS3*), and the other six likely affecting key T cell (*IL2RA, IL10, SIRPG, IKZF3, THEMIS*), thymus (*THEMIS*) and B cell development/functions (*IKZF3, IL10*) or in both immune and beta cells (*CTSH*) had stronger effects in the <7 group.

**Conclusions:** In newborn children with the greatest load of certain risk alleles, dysregulated response of immune and beta cells to environmental stresses such as virus infection, combine to cause a rapid loss of insulin production, driving down the age of type 1 diabetes diagnosis.

---

Type 1 diabetes is a multifactorial disease in which the insulin-producing beta cells of pancreatic islets are destroyed or rendered dysfunctional by an autoimmune process that often initiates in the first few months of life, causing a pre-diabetic, non-symptomatic state in approximately 0.4% of children ^1^. The actual diagnosis could happen many years after this prodromal phase, the joint environmental and genetic mechanisms of which remain ill defined, with the median age-at-diagnosis being around age 11 years. Even after diagnosis there often remains sufficient endogenous insulin production to lower the requisite levels of insulin treatment and reduce the later in life complications of early mortality, cardiovascular, kidney, eye and peripheral neuron disease ^2^. The exceptions to this are the children diagnosed with type 1 diabetes under the age five years in whom there is little insulin production shortly after diagnosis ^2,3^. This subgroup represents the largest unmet clinical challenge, since they suffer the greatest complications of the disease ^3^. Yet any intervention of type 1 diabetes autoimmunity in these young children must be as safe and precise as possible, modulating the causative molecules, cells, pathways and mechanisms. Hence we need to identify the specific mechanisms underlying early-diagnosed type 1 diabetes.

Recent evidence suggests that children diagnosed under age 7 years may have a different, more aggressive form of islet inflammation (insulitis), characterised by a B lymphocyte infiltrate coincident with a T cell insulitis (CD4^+^ and CD8^+^ T cells), than children aged 13 years and over, who have reduced B cell participation ^4^. In cases diagnosed between 7 and 12 years there is a mixture of islet infiltrate phenotypes, some with the “under 7” B cell infiltrate and others with “13 and over” phenotype. There is already evidence that autoantigen-presenting genes HLA class II and class I are associated with reduced age-at-diagnosis, which provides insight into the biology of this most beta-cell destructive form of the disease ^5–8^. More recently, a genome-wide association analysis of age-at-diagnosis of type 1 diabetes identified a locus on chromosome 6q22.33 that acts almost exclusively in type 1 diabetes cases diagnosed under age 5 years ^9^, encoding the protein tyrosine phosphatase receptor kappa (PTPRK) and the thymocyte-expressed molecule involved in selection (THEMIS) genes. However, this approach had to apply a stringent genome-wide multiple testing correction criterion (p <5×10^−8^) and informative, true signals were likely to have been missed. In the present study, we analysed the association of specified known type 1 diabetes risk regions, thereby reducing the multiple testing burden. In addition, a biological or phenotypic prior could provide greater sensitivity in the search for age-at-diagnosis-associated genes. The stratification of patients into age-at-diagnosis categories according to their pancreatic histology, as opposed to treating age-at-diagnosis as a continuous phenotype provides us with just this opportunity.

Here, we analysed type 1 diabetes-associated variants according to the proposed pancreatic infiltrate stratification of type 1 diabetes, namely the age-at-diagnosis groups, the under 7 group versus the 13 and over group. If type 1 diabetes has a particular pancreatic immunophenotype then it might be expected that it could have distinct genetic features, characterised by susceptibility genes with larger effects in the under 7 group. Moreover, the intermediate group, age-at-diagnosis 7-13 years, would have risk for these age-at-diagnosis-sensitive genes lying between the under 7’s and the 13’s and over. Six HLA haplotypes/alleles and seven non-HLA loci fulfil this risk profile informing the biology of the most aggressive form of type 1 diabetes, revealing a mixture of predisposition in both the beta cell and immune cell compartments.

## Research Design and Methods

### Study populations

Our dataset consists of 18,488 controls, 3,109 type 1 diabetes cases diagnosed at <7 years (‘<7 group’), 3,754 at ≥7 to <13 years (‘7-13 group’) and 1,724 at ≥13 years (‘≥13 group’). The majority of individuals are from the UK (Table 1), and comprises unrelated individuals, since related individuals were removed (Supplementary methods).

**Table 1:** Characteristics of individuals included in the analysis.

|  | **Controls** | **<7** | **7-13** | **≥13** |
| --- | --- | --- | --- | --- |
| N | 18488 | 3109 | 3754 | 1724 |
| Mean age-at-diagnosis | NA | 3.67 | 9.54 | 18.32 |
| Sex: Female | 9769 (52.9%) | 1493 (48.7%) | 1915 (52%) | 639 (42.9%) |
| Asia-Pacific | 927 (5%) | 22 (0.7%) | 23 (0.6%) | 36 (2.1%) |
| Central Europe | 1681 (9.1%) | 50 (1.6%) | 62 (1.7%) | 172 (10%) |
| Finland | 2824 (15.3%) | 100 (3.2%) | 154 (4.1%) | 438 (25.4%) |
| Northern Ireland | 479 (2.6%) | 222 (7.1%) | 248 (6.6%) | 35 (2%) |
| UK | 10593 (57.3%) | 2662 (85.6%) | 3206 (85.4%) | 923 (53.5%) |
| USA | 1984 (10.7%) | 53 (1.7%) | 61 (1.6%) | 120 (7%) |

### Loci studied

We examined eight HLA class II haplotypes and nine HLA class I classical alleles for their association with type 1 diabetes diagnosed within each age group, where the haplotypes and classical alleles were a subset of the most protective and susceptible haplotypes identified for type 1 diabetes to date ^10^ that we also found to be associated with type 1 diabetes in our analysis after conditioning on the other associated HLA haplotypes (logistic regression Wald test p<0.01). Supplementary Table 1 summarises which haplotypes and classical alleles were examined, how they were defined and whether they were common enough to include in our analysis, defined as having at least five individuals from each group carrying the classical allele/haplotype.

We also examined 55 loci outside the HLA, which have previously shown association with type 1 diabetes either in univariable analyses or via fine mapping (Supplementary Table 2). Each locus contains an ‘index’ variant, chosen to be the most strongly disease associated from a set of variants in linkage disequilibrium (LD) that constitute a single genetic signal. We have allocated locus names to each of these variants based on a candidate gene(s), but the named genes may not be causal for type 1 diabetes.

### Imputation

To impute classical HLA alleles, the HiBag ^11^ R Bioconductor package was used with the classifiers as calculated from the training dataset in the original publication. Alleles called with a probability of less than 0.5 were treated as missing. Some individuals were genotyped for a subset of their classical HLA alleles ^8^ and therefore accuracy of imputation was assessed at those classical alleles for a proportion of individuals.

When variants of interest failed genotype quality control (Supplementary methods), we imputed those variants plus 0.5 Mb surrounding regions using the IMPUTE2 ^12^ software; with the 1000 Genomes Project data as the reference haplotypes. We used this same imputation strategy to impute around index variants for fine mapping non-HLA regions found to be differentially associated between the <7 and ≥13 groups.

### Multinomial logistic regression

In order to examine whether or not there was heterogeneity in effect sizes between the <7 and ≥13 groups, we fitted two multinomial logistic regressions per locus, one allowing for different effect sizes for the genetic variant at that locus in the <7 and ≥13 groups and the other constraining the effect size for the genetic variant in the <7 and ≥13 groups to equal each other. As suggested in ^13^, we compared the likelihoods of these models using a likelihood ratio test, and if the model allowing different effect sizes fitted the data better than the model constraining the effect sizes to equal each other, the locus was considered heterogeneous in effect size between age-at-diagnosis groups. Both models included as covariates the ten largest principal components derived from the set of ImmunoChip variants passing quality control filters (Supplementary methods). This analysis was performed using the multinomRob R package ^14^.

For HLA loci, we additionally adjusted for other HLA haplotypes/alleles to account for the high levels of LD in the region. When examining the HLA class II haplotype effects other than DR3-DQ2/DR4-DQ8, we examined only individuals without the DR3-DQ2/DR4-DQ8 diplotype in order to remove any confounding effects of this diplotype. Supplementary Table 1 shows which individuals were included in each analysis, as well as the classical haplotypes/alleles adjusted for in each analysis.

### Sensitivity analyses – non-HLA results

To test stability of our results at non-HLA loci, we did three sensitivity analyses. Firstly, to exclude the possibility of spurious associations due to population structure in our data, we repeated the analysis using only individuals from the UK and Northern Ireland and adjusted for the five largest principal components derived from ImmunoChip data in these individuals only. Additionally, to test sensitivity of our results to age-strata thresholds, we performed the same analysis but instead compared individuals diagnosed at <6 years to the ≥13 group and also individuals diagnosed at <5 years compared to the ≥13 group.

We declared a locus differentially-associated if the heterogeneity p-value was associated to a False Discovery Rate (FDR) of <0.1 (Supplementary methods). To explore whether there were more age-at-diagnosis associated variants which we could not detect in the present analysis due to a lack of statistical power, we examined all loci which did not reach the association threshold (FDR<0.1) by counting how many loci had the largest effect in the <7 group, the intermediate effect in the 7-13 group and the smallest effect in the ≥13 group and compared this to the expected frequency of this ordering using a binomial test (Supplementary methods).

### Fine mapping

For the non-HLA loci with the evidence of heterogeneity in effect size between age-at-diagnosis groups (FDR<0.1), we fine mapped a 0.5 Mb region around the index variant to identify a list of potentially causal variants for type 1 diabetes diagnosed at <7 years. Analysis was limited to individuals from the UK and Northern Ireland, amounting to 2,884 cases diagnosed at <7 years and 11,072 controls, in order to examine a homogeneous population, as fine mapping is sensitive to differences in LD structure between ancestrally divergent groups. We used the GUESSFM software, ^15^ which carries out a Bayesian variable selection stochastic search to identify the combinations of variants constituting separate genetic susceptibility to type 1 diabetes (Supplementary methods).

### Colocalisation analyses

For regions with evidence of heterogeneity in effect size between age-at-diagnosis groups and also those that had high (>0.8) posterior probability of a single causal variant in the region, we conducted colocalisation analyses with expression quantitative trait loci (eQTL) associations in whole blood from a dataset of over 30,000 individuals ^16^. This allows assessment of the genes the variants are most likely regulating and what direction the effects are on gene transcription and disease risk. The coloc R package was used to carry out this analysis ^17^ (Supplementary methods).

### Heritability estimates by age-at-diagnosis group

We compared chip heritability estimates between type 1 diabetes cases diagnosed at <7 years to those diagnosed at 7-13 years or ≥13 years. We used the GCTA software (https://cnsgenomics.com/software/gcta/#Overview) to fit linear mixed models and estimate heritability for each age-at-diagnosis group with a shared set of controls. We estimated heritability, h_g_^2^, on the liability scale, as derived in ^18^. We adjusted for the ten largest principal components as in the multinomial regression analysis and included individuals from all ancestry backgrounds in the analysis. We repeated the analysis but excluded the entirety of the HLA region. Chip heritability using the ImmunoChip is likely to not reflect the ‘true’ heritability, since there are large regions of the genome without any genotype data available, with approximately ~117,000 included after quality control. The prevalence of type 1 diabetes between age-at-diagnosis groups may vary; we assume a prevalence of 0.4% in all age groups in the primary analysis but test stability of the estimates by performing sensitivity analyses where the prevalence in the <7 group was higher at 0.5% and also examining heritability in the 7-13 and ≥13 group with an assumed prevalence of both 0.2% and 0.3%.

### Code availability

The scripts used to analyse these data are available at https://github.com/jinshaw16/AAD_t1d.

## Results

### Multinomial logistic regression: HLA

We found six HLA variables to be differentially-associated between the <7 and ≥13 group (FDR<0.1). The strongest susceptible class II effect was for the DR3-DQ2/DR4-DQ8 diplotype, whilst the protective DRB1*15:01-DQB1*06:02 and DRB1*07:01-DQB1*03:03 haplotypes showed greater protection from type 1 diabetes in the <7 group compared to the ≥13 group. Class I alleles A*24:02 and B39*06 showed more susceptibility to type 1 diabetes in the <7 compared to and ≥13 group (Figure 1).

**Figure 1:**
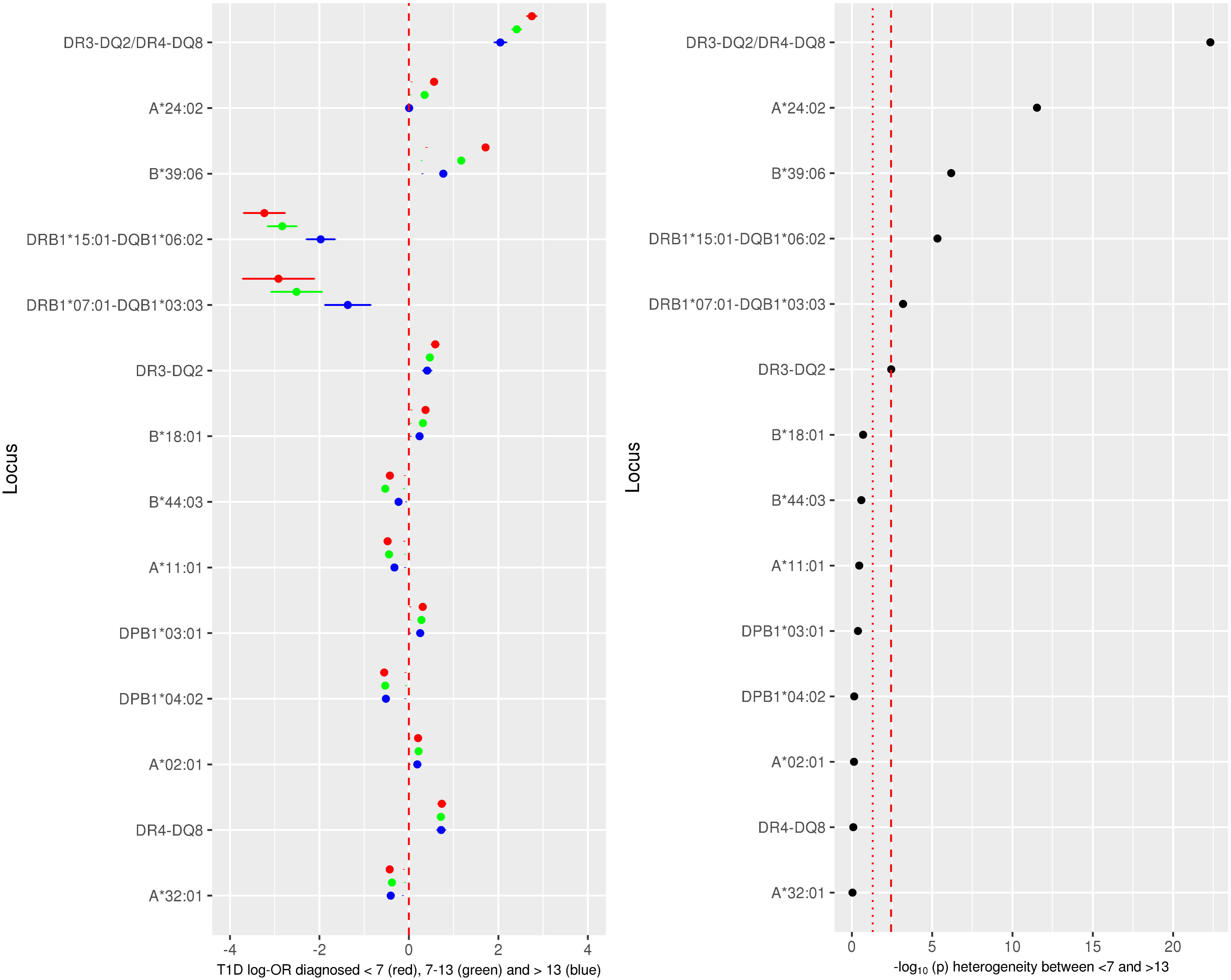
Classical HLA haplotype/alleles association with type 1 diabetes diagnosed at <7 years old (red circle; mean log-odds ratio age-at-diagnosis +/- 95%CI), 7-13 years old (green circle; mean log-odds ratio age-at-diagnosis 7-13+/- 95%CI) and ≥13 years old (blue circle mean log-odds ratio age-at-diagnosis >13+/- 95%CI), from a multinomial logistic regression. Left panel shows the log-odds ratios with a dashed red line showing a log-odds ratio of 0. The right panel shows the association statistics from a likelihood ratio test comparing a multinomial logistic regression constraining the log-odds ratios from the <7 and ≥13 groups to be equal compared to an unconstrained model. Red dotted line shows nominal significance in heterogeneity (p<0.05), red dashed line show Bonferroni-corrected significance in heterogeneity.

Comparison of imputed classical 4 digit HLA alleles with directly genotyped 4 digit HLA alleles showed concordance of over 91% for each gene examined (Supplementary Figure 1).

### Multinomial logistic regression: non-HLA regions

Outside the HLA, nine regions were differentially-associated between the <7 and ≥13 group (FDR<0.1), near Cathepsin H (CTSH), GLIS family zinc finger 3 (GLIS3), Ikaros family zinc finger 3 (IKZF3), the third index variant at interleukin 2 receptor alpha (IL-2RA), Chymotrypsinogen B1 (CTRB1), Calmodulin-Regulated Spectrin-Associated Protein 2 (CAMSAP2), interleukin 10 (IL-10), Signal Regulatory Protein Gamma (SIRPG) and THEMIS (Figure 2). Two of these genes (*CTSH* and *GLIS3*) survived Bonferroni correction (p<0.05/55=0.00091). At each locus associated with FDR<0.1, the 7-13 group had a larger effect size than the ≥13 group and smaller than the <7 group. Given the ≥13 group comprises just 1,724 individuals, it is probable that with increased sample size and hence statistical power, other type 1 diabetes risk loci might reach statistical significance with regards to heterogeneity (Supplementary Figure 2). Of the 46 variants not satisfying an FDR<0.1, 20 have the strongest signal in <7s, weakest in ≥13s and intermediate in 7s-13s, compared to eight occurrences in that order expected by chance (p=4.27×10^−6^, binomial test), suggesting the presence of substantial additional signal in variants that we are not able to declare show evidence individually.

**Figure 2:**
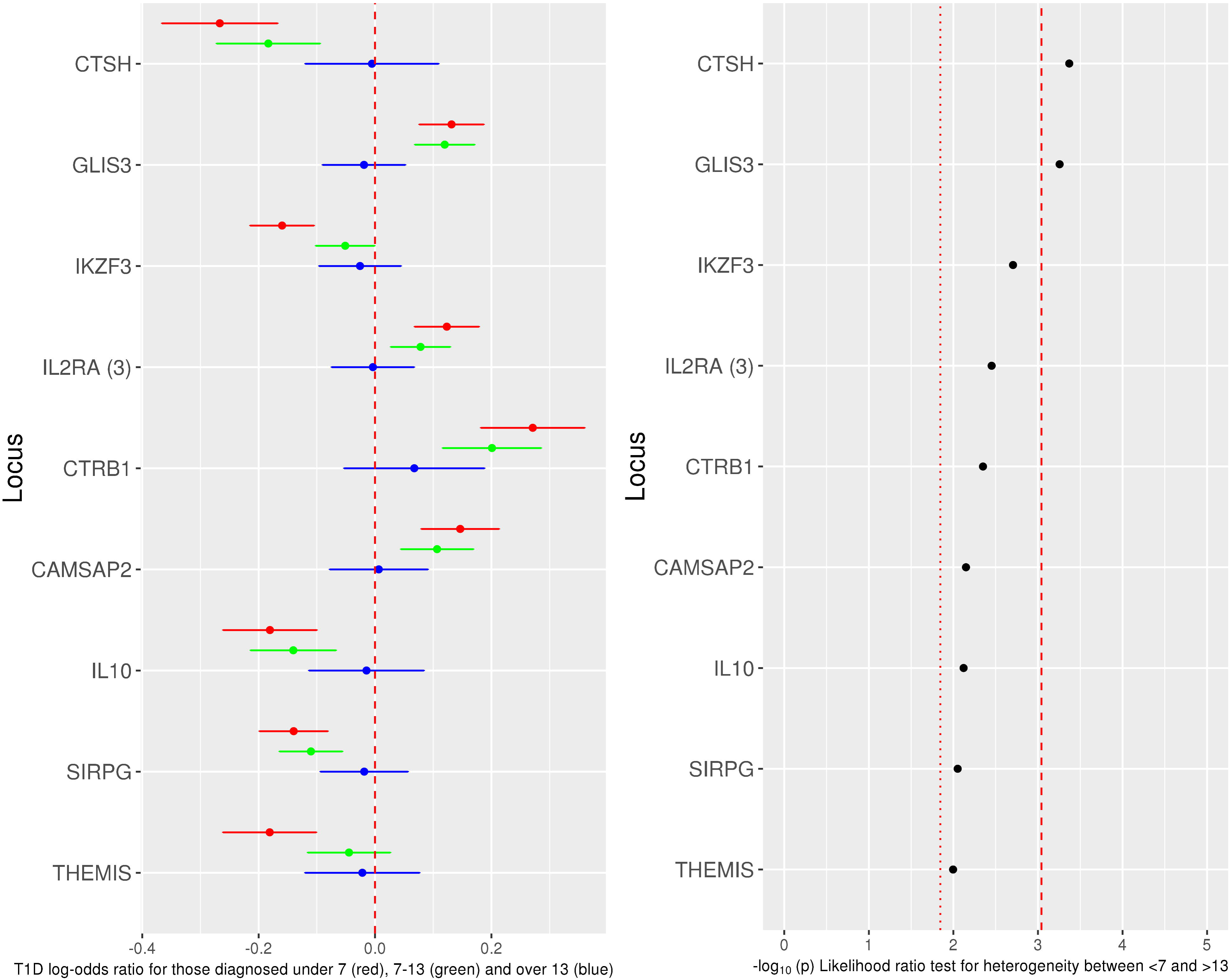
Non-HLA type 1 diabetes associated variants, showing log-odds ratios for those diagnosed at <7 years old (red circle; mean log-odds ratio age-at-diagnosis +/- 95%CI), 7-13 years old (green circle; mean log-odds ratio age-at-diagnosis 7-13+/- 95%CI) and ≥13 years old (blue circle mean log-odds ratio age-at-diagnosis ≥13+/- 95%CI), from a multinomial logistic regression. Left panel shows the log-odds ratios with a dashed red line showing a log-odds ratio of 0. The right panel shows the association statistics from a likelihood ratio test comparing a multinomial logistic regression constraining the log-odds ratios from the <7 and ≥13 groups to be equal compared to an unconstrained model. Showing only loci with a false discovery rate of less than 0.1. Red dotted line shows threshold for false discovery rate of <0.1, red dashed line shows threshold for Bonferroni-corrected heterogeneity.

### Stability of non-HLA results

In the UK-specific sensitivity analysis, six of the nine loci declared heterogeneous from the primary analysis were heterogeneous between the <7 and ≥13 group (FDR<0.1), two of the loci showed no heterogeneity in effect size (*CTRB1* p=0.310 and *CAMSAP2* p*=*0.578) and were thus removed from our set of differentially-associated regions and the remaining locus, *IL10*, had a p-value of 0.08, which we considered differentially associated between the <7 and ≥13 groups, given the decrease in statistical power in this sensitivity analysis (Supplementary Figure 3). When changing the threshold for the early-diagnosed group to <6 and <5, all seven associated loci from the primary analysis and UK-specific analysis were heterogeneous (FDR<0.1) (Supplementary Figures 4 and 5).

Minor allele frequency plots by age-at-diagnosis for the seven differentially-associated loci are shown in Supplementary Figures 6-12, whilst Supplementary Tables 3 and 4 summarise the most likely causal genes at these loci.

### Fine mapping

We fine mapped the seven differentially-associated loci. Five regions had high posterior probability of a single causal variant in the region (>0.8), whilst *SIRPG* (0.69) and *IL2RA* (5.41×10^−7^) had a higher posterior probability of having more than one causal variant in the region. Variants prioritised in the GUESSFM analysis for each region examined are listed in Supplementary Tables 5-11, though it is possible that variants excluded due to low imputation quality that are in LD with these variants could also be causal.

Three of the five regions fine-mapped with high posterior probability of one causal variant in the region showed evidence of colocalisation with whole-blood eQTLs. The *CTSH* locus showed evidence of colocalisation with a *CTSH* eQTL (posterior probability of colocalisation=0.998); the susceptibility allele for type 1 diabetes is associated with more expression of *CTSH*. The *IKZF3* locus fine mapping results prioritises an LD block containing 99 variants, all of which could be causal, which also effects expression of at least three genes (p<5 × 10^−150^), with evidence of colocalisation of disease signal and whole-blood eQTL for all of *IKZF3, GSDMB* and *ORMDL3* (posterior probability of colocalisation for type 1 diabetes and eQTL with *IKZF3*=0.983, *GSDMB*=0.974 and *ORMDL3*=0.974); the most likely causal variants decrease type 1 diabetes risk and *IKZF3* expression and also increase expression of *GSDMB* and *ORMDL3*. Finally, the most likely causal variants in the *THEMIS* region colocalise with a *THEMIS* whole-blood eQTL (posterior probability of colocalisation=0.953); the susceptible type 1 diabetes alleles are associated with increased *THEMIS* expression in whole blood (Figure 3).

**Figure 3:**
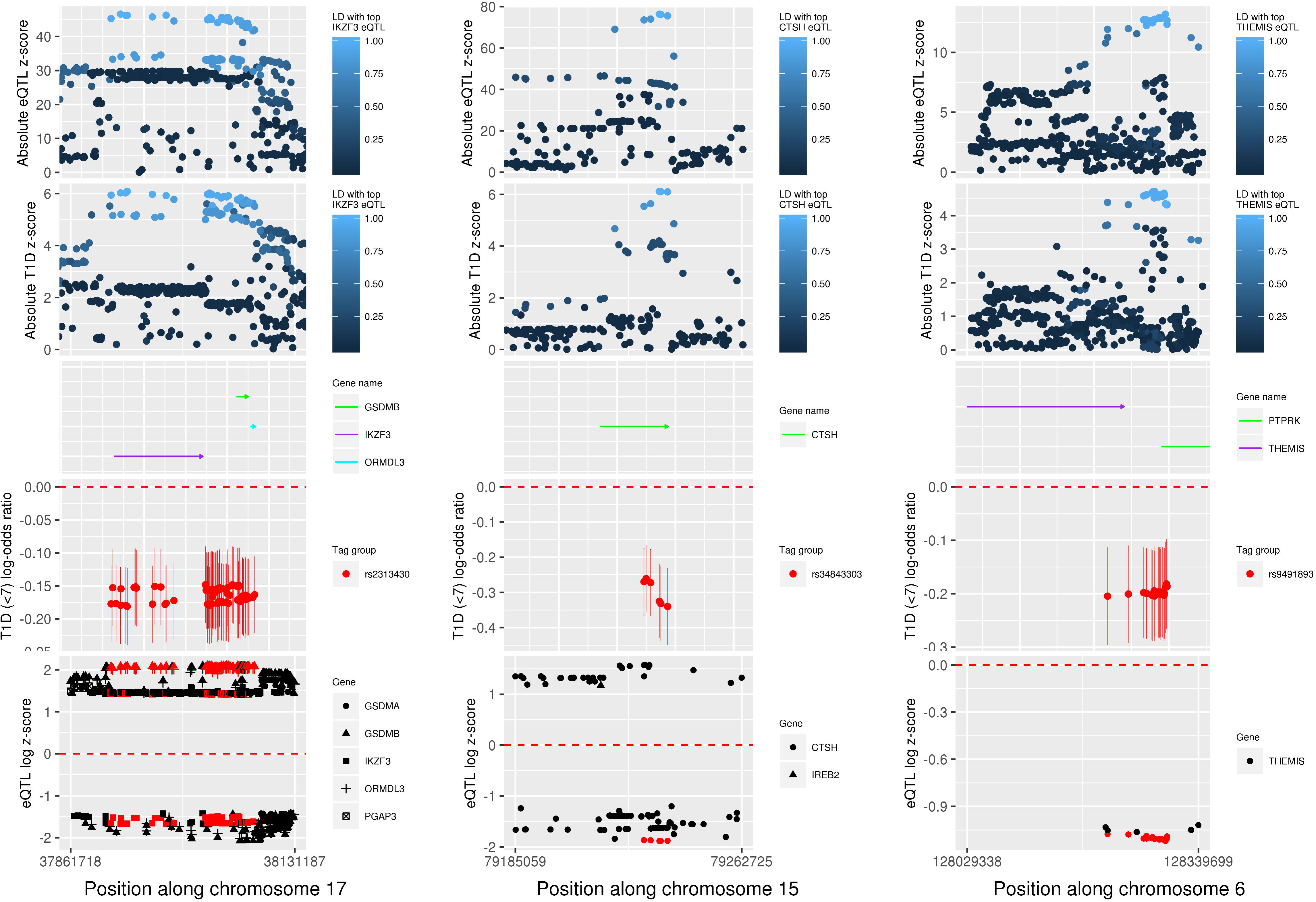
Results from colocalisation and fine mapping in the *IKZF3* region (left) *CTSH* region (centre) and THEMIS region (right). Analyses include individuals from the UK and Northern Ireland and only controls and cases diagnosed at <7 years. First panel: association absolute z scores from whole blood eQTL study examining variant effects on *CTSH* (left), *IKZF3* (centre) and *THEMIS* (right) mRNA levels, coloured by LD r^2^ to the most strongly associated variant with the respective mRNA expression. Second panel: association absolute z scores from logistic regression examining variant associations with type 1 diabetes risk at < 7 years, coloured by LD r^2^ to the most strongly associated variant with *CTSH* (left), *IKZF3* (centre) and *THEMIS* (right) mRNA. Third panel: Gene positions (genome build 37), with arrows indicating direction of transcription. Fourth panel, univariable early-diagnosed type 1 diabetes log-odds ratios and 95% confidence intervals for each of the most likely causally associated variants as prioritised by GUESSFM. Fifth panel: log_e_(absolute eQTL z score) if z score>0 and −log_e_(absolute eQTL z score) if z score<0, so direction of effect can be compared, including only eQTLs with a p value of <5×10^−150^ (left),<5×10^−50^ (centre) and <5×10^−25^ (right). The symbols are coloured red if contained in the set of most likely causal variants, as produced by GUESSFM and the shape corresponds to the gene that the variant is effecting transcription of.

There was no evidence of colocalisation between disease risk variants and whole blood eQTL for any genes in the region of *GLIS3* or *IL10* (posterior probability of colocalisation<0.01), suggesting the variants in these regions might be acting in other cell types or activation states to alter type 1 diabetes risk (Supplementary Figures 13-16).

### Heritability estimates by age-at-diagnosis group

We found the chip heritability estimate on the liability scale to be highest in the <7 group, intermediate in the 7-13 group and lowest in the ≥13 group (<7 h_g_^2^=0.366, 7-13 h_g_^2^=0.301, ≥13 h_g_^2^=0.236). This trend remained when altering the assumed disease prevalence by age-at-diagnosis group and when excluding the HLA region (Supplementary Table 12).

## Conclusions

The stratification of patients by age-at-diagnosis according to islet phenotypes has provided a rich source of genes, molecules and pathways with greater effects in children diagnosed with type 1 diabetes under age 7 years. We expected to see strong differential associations with the HLA class II haplotypes, in particular the heterozygous diplotype DR3-DQ2/DR4-DQ8, as well as HLA class I alleles, A*24:02 and B*39:06 ^5–8^. Here, we show for the first time that the protective HLA class II haplotypes DRB1*15:01-DQB1*06:02 and DRB1*07:01-DQB1*03:03 are less prevalent amongst individuals diagnosed at <7 years compared with controls and those diagnosed at ≥13 years. Therefore, the earliest and most aggressive phenotypic subtype of type 1 diabetes results primarily from carriage of high risk alleles of the HLA class II and I genes, which probably act at one or more of four levels: (i) altering the T cell receptor repertoire in favour of anti-islet antigen reactivity, for example preproinsulin, and/or reducing the protective repertoire of T regulatory cells; (ii) providing a strong autoantigen presentation environment in the islets and pancreatic draining lymph nodes enabling the infiltration and cytolytic activity of CD8^+^ T cells but also by disrupting B cell anergy ^19^ permitting binding and presentation of autoantigen to provide potent help to T cells in a self-reinforcing spiral of autoreactivity; (iii) affecting the immune response to the viral infections that are involved in the disease; (iv) affecting how the gut microbiome develops in early life, a system that is known to affect type 1 diabetes susceptibility ^20^.

In addition to the HLA heterogeneity, we obtained robust evidence of differences in effect size between the age-at-diagnosis groups at seven non-HLA loci. Of these loci, one plausible candidate gene, *GLIS3*, most likely perturbs disease risk in the islet beta cells, given the expression levels in the pancreas, lack of expression in immune cells, colocalisation with type 2 diabetes risk variants ^21^ and lack of association with other autoimmune diseases (https://genetics.opentargets.org). Genes in two of the loci, *CTSH* and *IKZF3*, could act in the islets or elsewhere, whilst all of the other candidate causal genes (*IL2RA, IL10, SIRPG, THEMIS, IKZF3/ORMDL3/GSDMB* and *CTSH*) have known functions in T and/or B cell biology (Supplementary Table 4). This implies that in addition to HLA-susceptibility, risk of type 1 diabetes in the very young is also impacted by particular malfunctions in the infiltrating T and B cells, leading to increased risk of autoreactivity, resulting in a perfect storm of immune infiltration, antigen recognition and a rapid destruction of beta cells.

Of the seven non-HLA risk regions with the strongest evidence of heterogeneity between age-at-diagnosis groups, we focus on the *CTSH, IKZF3* and *THEMIS* loci, which colocalise with whole blood eQTLs. The candidate type 1 diabetes risk variants at the *CTSH* locus, for example the C allele at rs2289702 (C>T), are associated with increased expression of CTSH mRNA in multiple cell types and tissues (Supplementary Table 4). The locus has previously been implicated in type 1 diabetes aetiology by altering sensitivity of beta cells to apoptosis ^22^, where rs3825932 (C>T) was investigated, which is in low LD (r^2^=0.26) with the disease-associated variant reported here. However, the type 1 diabetes risk allele counter-intuitively resulted in protection from beta-cell apoptosis, thus, beta-cell apoptosis may not be the primary mechanism underlying disease aetiology in this region. *CTSH* functions as an endopeptidase and can cleave the N-terminus of the Toll-like receptor 3 (TLR3) protein, increasing its functionality ^23^. Given TLR3 is expressed in islets ^24^, it is possible that the increase in CTSH expression associated with the type 1 diabetes susceptibility allele results in increased TLR3 N-terminus cleavage, heightened responses to viral infections and increased release of type 1 interferon (IFN). This may increase baseline risk of type 1 diabetes and specifically the risk of early-diagnosed type 1 diabetes in individuals carrying this allele, since viral infections are more frequent in childhood. There is mounting evidence that enteroviral infections predispose to type 1 diabetes: a type 1 IFN transcriptional signature precedes anti-islet autoantibody appearance in children ^25^; and another receptor for viral RNA, MDA5 encoded by *IFIH1* is a proven type 1 diabetes susceptibility gene with its higher IFN-inducing activity increasing risk of the disease ^26^. Exposure of beta cells to type 1 IFN greatly increases their HLA class I expression and susceptibility to CD8^+^ cytotoxic killing, and heightened class I expression on beta cells is a hallmark phenotype of the type 1 diabetes pancreas ^27^.

The region containing *IKZF3* has a large LD block, which is associated with multiple diseases, including asthma and paediatric asthma ^28^, ^29^. However, the direction of effect of the risk variant is opposite in asthma to all associated autoimmune diseases, where the C allele at a variant within the haplotype, rs921649 (C>T), increases susceptibility to autoimmunity, whereas the C allele is protective for asthma ^28^. The three genes most strongly associated with the haplotype are expressed in lymphocytes and are up-(*IKZF3*) or down-regulated (*ORMDL3, GSDMB*) (https://dice-database.org/) following activation, with good biological candidacy for altering disease risk. *IKZF3* is a transcriptional repressor with a key role in B-cell activation and differentiation ^30^ and T cell differentiation ^31^. *ORMDL3* is a central regulator of sphingolipid biosynthesis ^32^ and has also been proposed to negatively regulate store-operated calcium, lymphocyte activation and cytokine production ^28,33^, while *GSDMB* can act as a pyroptotic protein ^34^. Therefore one or more of these genes may be causal for type 1 diabetes risk. Pertinent to the increased frequency of B-cell infiltration in the islets of the <7 group, there is evidence that carriers of the type 1 diabetes risk allele have decreased anergic high affinity insulin-binding B cells in circulating blood, implying some of this population may have relocated to the pancreas ^19^. The colocalisation between the type 1 diabetes disease association and a *THEMIS* whole blood eQTL points towards *THEMIS* being the more likely causal gene in this region. The T allele at one of the candidate causal variants, rs13204742 (G>T) is protective for type 1 diabetes with early diagnosis and coeliac disease ^35^, but susceptible for irritable bowel disease ^36^. *THEMIS*, expressed in thymocytes and circulating T cells (Supplementary Table 4), is a key signalling molecule for T cell development and survival and may have different function in the thymus and periphery ^37^. In the thymus, *THEMIS* sets signalling thresholds at the double positive stage of thymocyte development and influences selection of T cells. Deletion of *THEMIS* reduces transition of double positive to single positive thymocytes ^38,39^. Our results suggest that an increase in *THEMIS* expression leads to increased risk of type 1 diabetes diagnosed at <7 years. If we assume the whole blood eQTL with the disease risk variants is mirrored in the thymus, we hypothesise that the increased *THEMIS* expression would alter the threshold for positive selection and increase the probability of autoreactive T cells entering circulation.

The increased heritability of early-diagnosed compared to later-diagnosed type 1 diabetes should be interpreted with caution given the ImmunoChip design is primarily focussed on immune regions and designed to capture regions of interest in autoimmune diseases. It is possible that those individuals diagnosed with type 1 diabetes later in life have a different profile of susceptibility regions that are not captured on the ImmunoChip array that we have not had statistical power to detect in previous GWAS studies. Nevertheless, it is interesting to note that amongst the susceptibility regions for type 1 diabetes discovered to date, there appears to be a higher heritability in those diagnosed at a young age compared to at 13 years old or older, consistent with increasing age accompanying greater risk and length of exposure to environmental type 1 diabetes causal factors.

Our genetic results imply a dynamic, fully integrated pathogenic collaboration between the immune system, the beta cells and viral infection in the initiation and rapid development of extreme insulin-deficiency starting in the first few weeks and months of life in those that carry the heaviest load of age-at-diagnosis alleles. Combinations of modulators of these pathways could be an effective way of preventing the cessation of endogenous insulin-production.

## Acknowledgments

We gratefully acknowledge all participants for allowing the analysis of their genotypic and phenotypic data.

## Author contributions

JRJI carried out the analyses and drafted the main body of the manuscript. AJC provided assistance with the manuscript writing and carried out a thorough literature search for age-at-diagnosis associated regions. DJMC provided statistical support and guidance for JRJI as well as critically reviewing the manuscript. LSW provided critical feedback for the manuscript and immunological support and context. JAT supervised the work, proposed the research hypothesis and critically evaluated the manuscript.

JAT is the guarantor for this manuscript.

No authors have any conflict of interests to declare.

This work was funded by the Juvenile Diabetes Research Foundation (JDRF) (9-2011-253, 5-SRA-2015-130-A-N) and Wellcome (091157, 107212) to the Diabetes and Inflammation Laboratory, University of Oxford.

We use data generated by the Wellcome Trust Case Control Consortium (076113). The Northern Irish Genetic Resource Investigating Diabetes (GRID), Tyypin 1 Diabetekseen Sairastuneita Perheenjäsenineen□(IDDMGEN), Tyypin 1 Diabetekseen Genetiikka (T1DGEN) and Warren cohorts were genotyped using the Type 1 Diabetes Genetics Consortium (T1DGC) grants from the National Institute of Diabetes and Digestive and Kidney Diseases (NIDDK), the National Institute of Allergy and Infectious Diseases (NIAID), the National Human Genome Research Institute (NHGRI), the National Institute of Child Health and Human Development (NICHD) and the JDRF (U01 DK062418, JDRF 9-2011-530). The research was supported by the Wellcome Trust Core Award Grant Number 203141/Z/16/Z with additional support from the NIHR Oxford BRC.

---

**Supplementary Figure 1:**
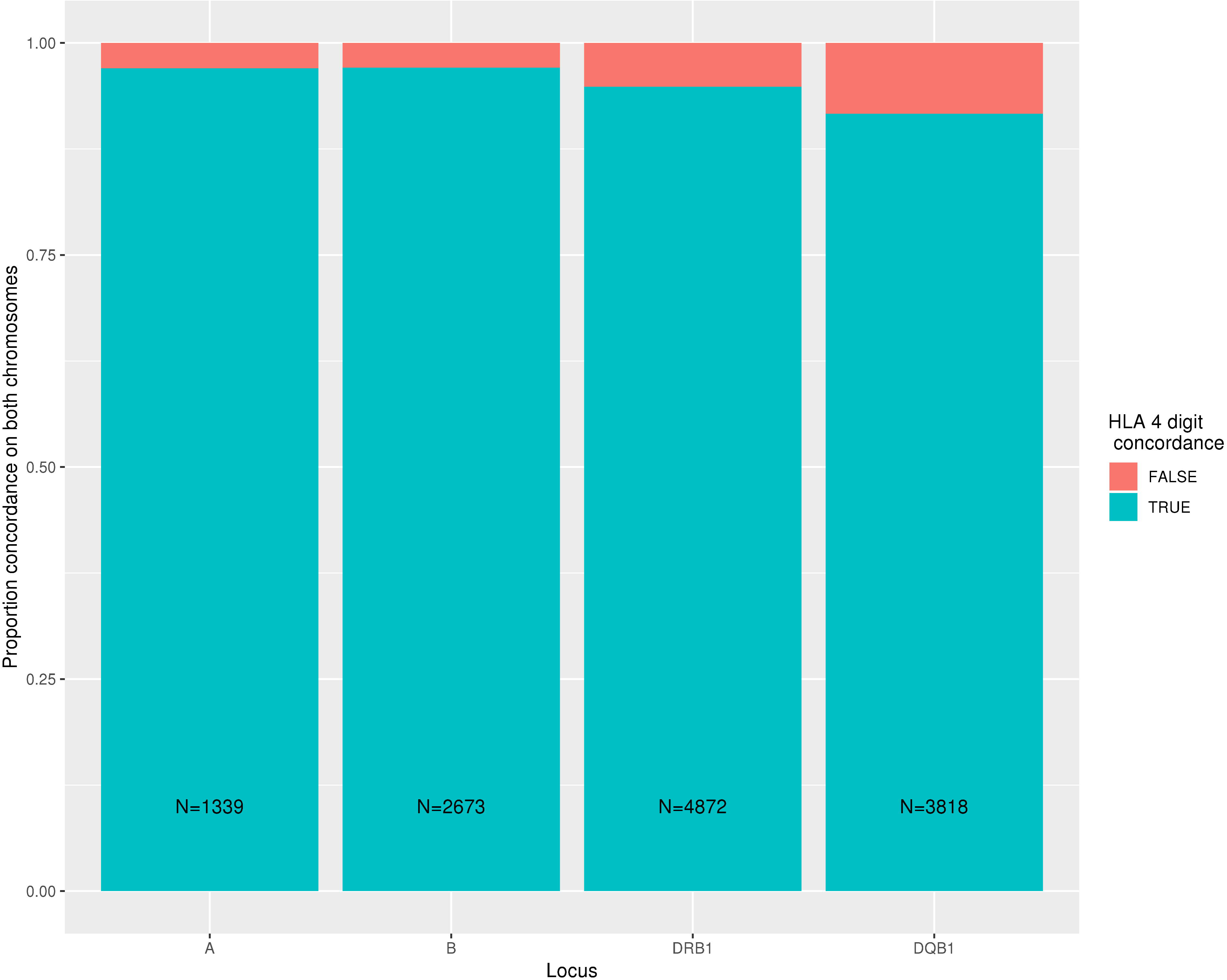
Concordance of HiBag imputed versus directly genotyped classical HLA alleles. Concordance is defined as identical 4 digit HLA classical allele at both chromosomes.

**Supplementary Figure 2:**
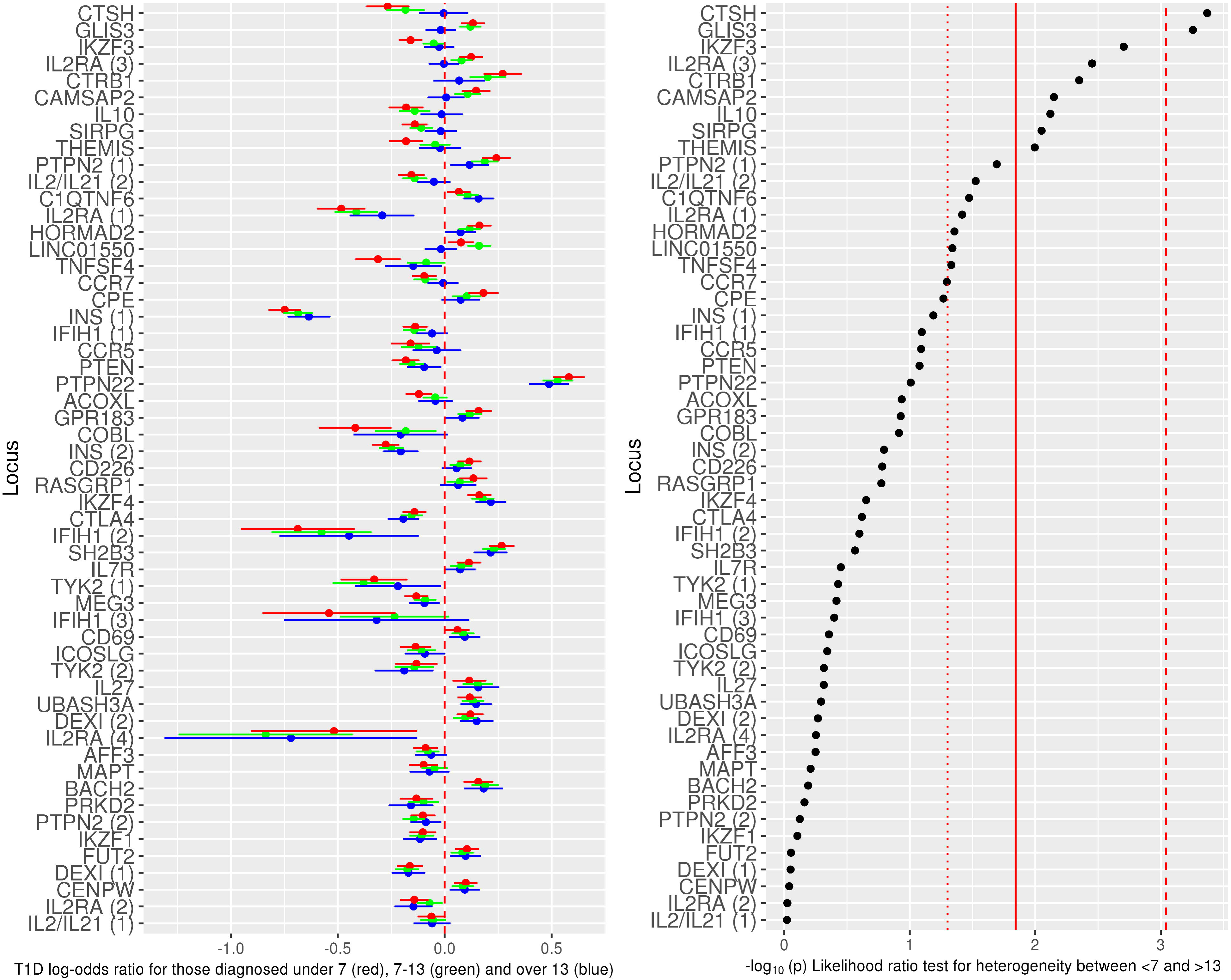
All 55 non-HLA type 1 diabetes associated variants, showing log-odds ratios for those diagnosed at <7 years old (red circle; mean log-odds ratio age-at-diagnosis +/- 95%CI), 7-13 years old (green circle; mean log-odds ratio age-at-diagnosis 7-13+/- 95%CI) and ≥13 years old (blue circle mean log-odds ratio age-at-diagnosis ≥13+/- 95%CI), from a multinomial logistic regression. Left panel shows the log-odds ratios with a dashed red line showing a log-odds ratio of 0. The right panel shows the association statistics from a likelihood ratio test comparing a multinomial logistic regression constraining the log-odds ratios from the <7 and ≥13 groups to be equal compared to an unconstrained model. Red dotted line shows threshold for nominally significant heterogeneity between groups (p<0.05), red solid line shows threshold for false discovery rate of <0.1, red dashed line shows threshold for Bonferroni-corrected significant heterogeneity.

**Supplementary Figure 3:**
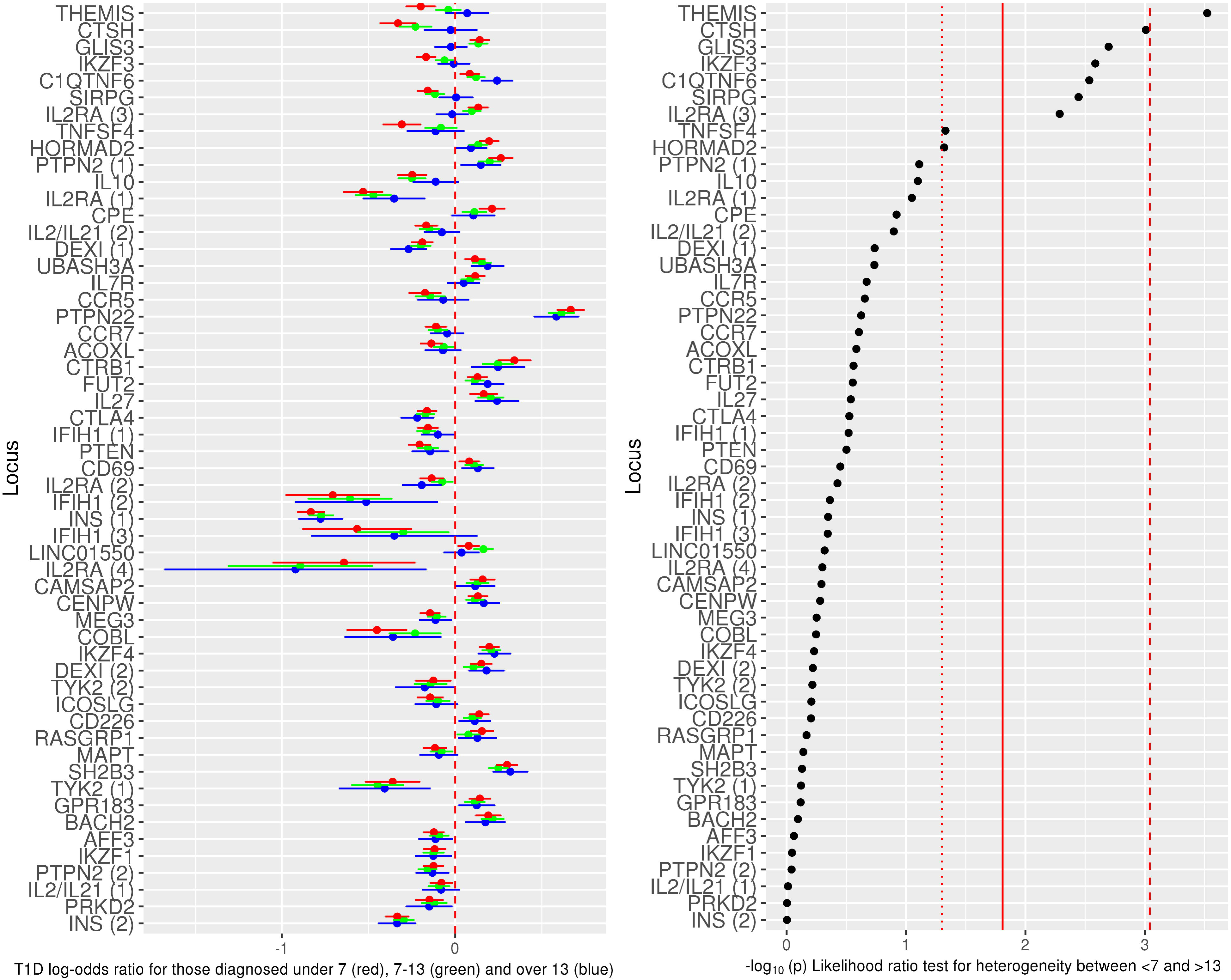
All 55 non-HLA type 1 diabetes associated variants examined using only individuals from the UK or Northern Ireland, showing log-odds ratios for those diagnosed at <7 years old (red circle; mean log-odds ratio age-at-diagnosis +/- 95%CI), 7-13 years old (green circle; mean log-odds ratio age-at-diagnosis 7-13+/- 95%CI) and ≥13 years old (blue circle mean log-odds ratio age-at-diagnosis >13+/- 95%CI), from a multinomial logistic regression. Left panel shows the log-odds ratios with a dashed red line showing a log-odds ratio of 0. The right panel shows the association statistics from a likelihood ratio test comparing a multinomial logistic regression constraining the log-odds ratios from the <7 and ≥13 groups to be equal compared to an unconstrained model. Red dotted line shows threshold for nominally significant heterogeneity between groups (p<0.05), red solid line shows threshold for false discovery rate of <0.1, red dashed line shows threshold for Bonferroni-corrected significant heterogeneity.

**Supplementary Figure 4:**
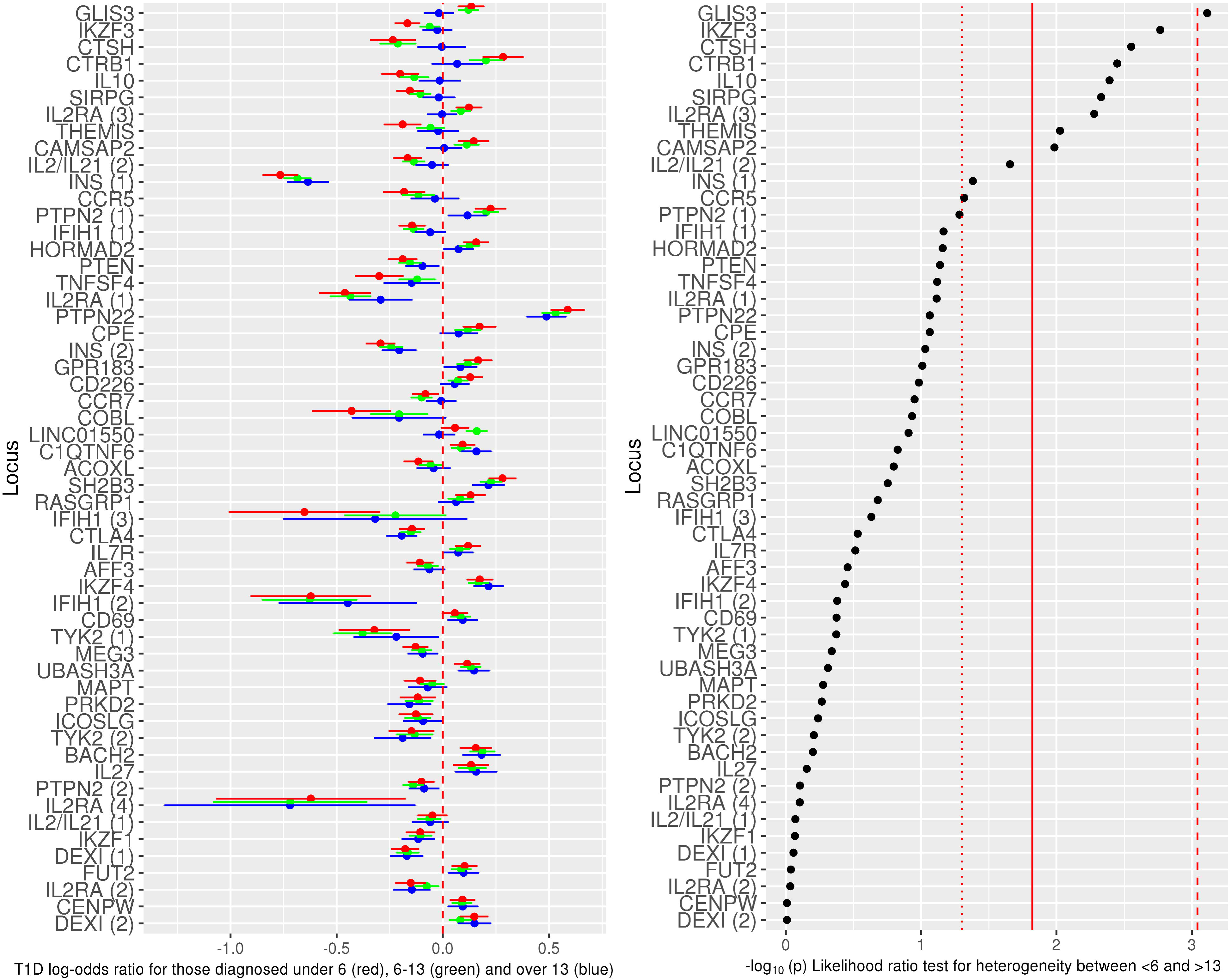
All 55 non-HLA type 1 diabetes associated variants, showing log-odds ratios for those diagnosed at <6 years old (red circle; mean log-odds ratio age-at-diagnosis +/- 95%CI), 6-13 years old (green circle; mean log-odds ratio age-at-diagnosis 6-13+/- 95%CI) and ≥13 years old (blue circle mean log-odds ratio age-at-diagnosis ≥13+/- 95%CI) from a multinomial logistic regression. Left panel shows the log-odds ratios with a dashed red line showing a log-odds ratio of 0. The right panel shows the association statistics from a likelihood ratio test comparing a multinomial logistic regression constraining the log-odds ratios from the <6 and ≥13 groups to be equal compared to an unconstrained model. Red dotted line shows threshold for nominally significant heterogeneity between groups (p<0.05), red solid line shows threshold for false discovery rate of <0.1, red dashed line shows threshold for Bonferroni-corrected significant heterogeneity.

**Supplementary Figure 5:**
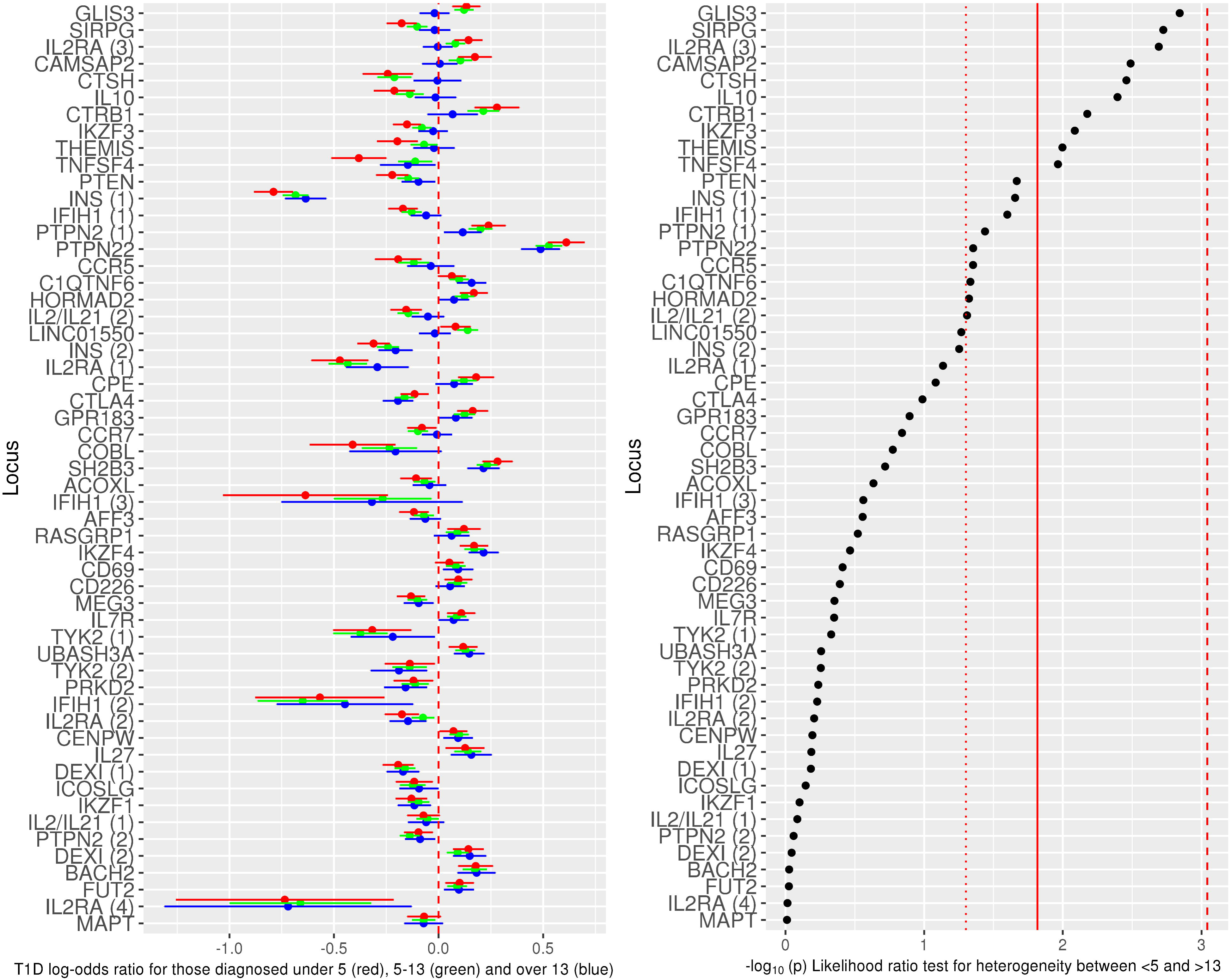
All 55 non-HLA type 1 diabetes associated variants, showing log-odds ratios for those diagnosed at <5 years old (red circle; mean log-odds ratio age-at-diagnosis +/- 95%CI), 5-13 years old (green circle; mean log-odds ratio age-at-diagnosis 5-13+/- 95%CI) and ≥13 years old (blue circle mean log-odds ratio age-at-diagnosis ≥13+/- 95%CI) from a multinomial logistic regression. Left panel shows the log-odds ratios with a dashed red line showing a log-odds ratio of 0. The right panel shows the association statistics from a likelihood ratio test comparing a multinomial logistic regression constraining the log-odds ratios from the <7 and ≥13 groups to be equal compared to an unconstrained model. Red dotted line shows threshold for nominally significant heterogeneity between groups (p<0.05), red solid line shows threshold for false discovery rate of <0.1, red dashed line shows threshold for Bonferroni-corrected significant heterogeneity.

**Supplementary Figure 6:**
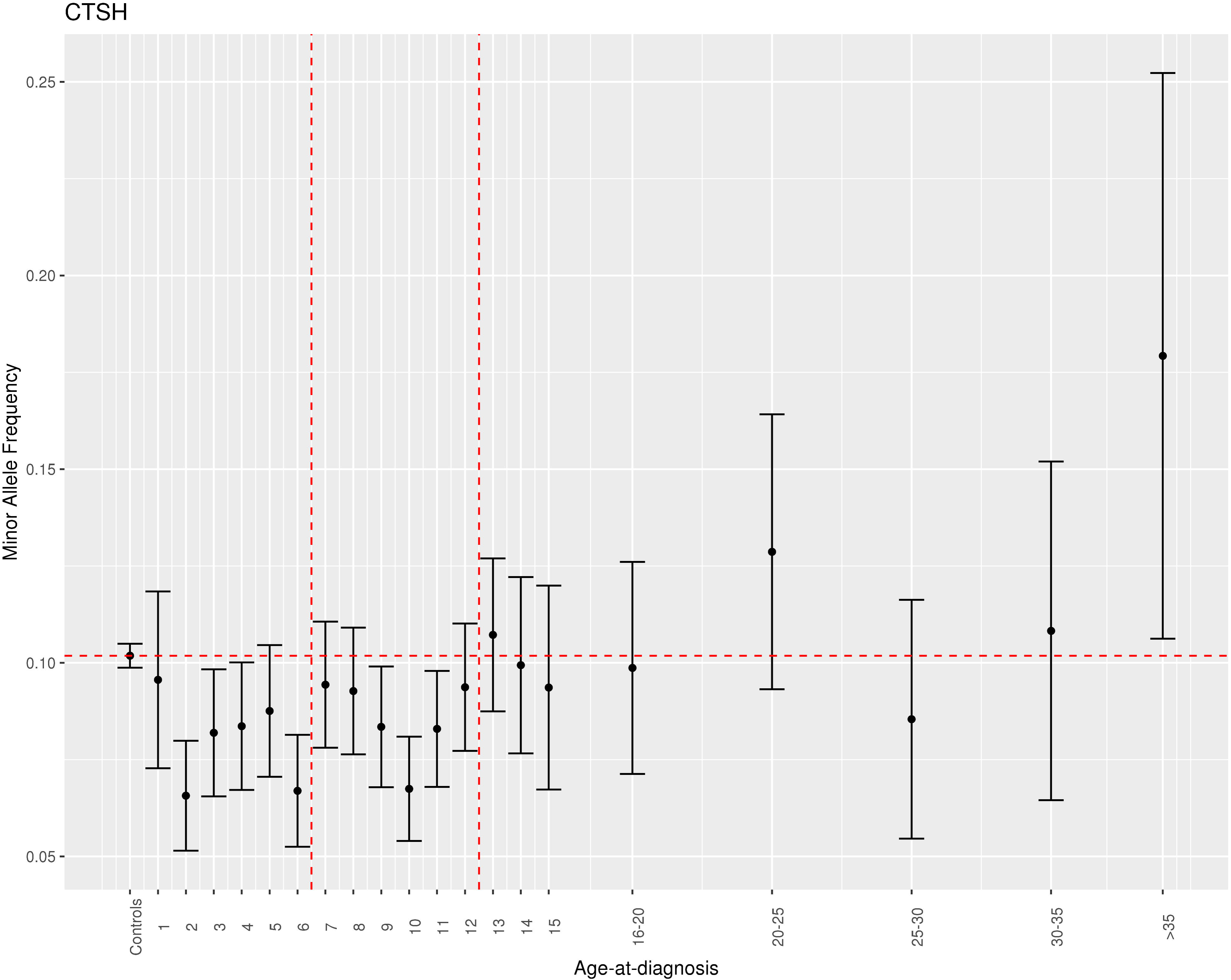
Minor allele frequency of the index variant near the *CTSH* gene for controls and individuals diagnosed at various ages.

**Supplementary Figure 7:**
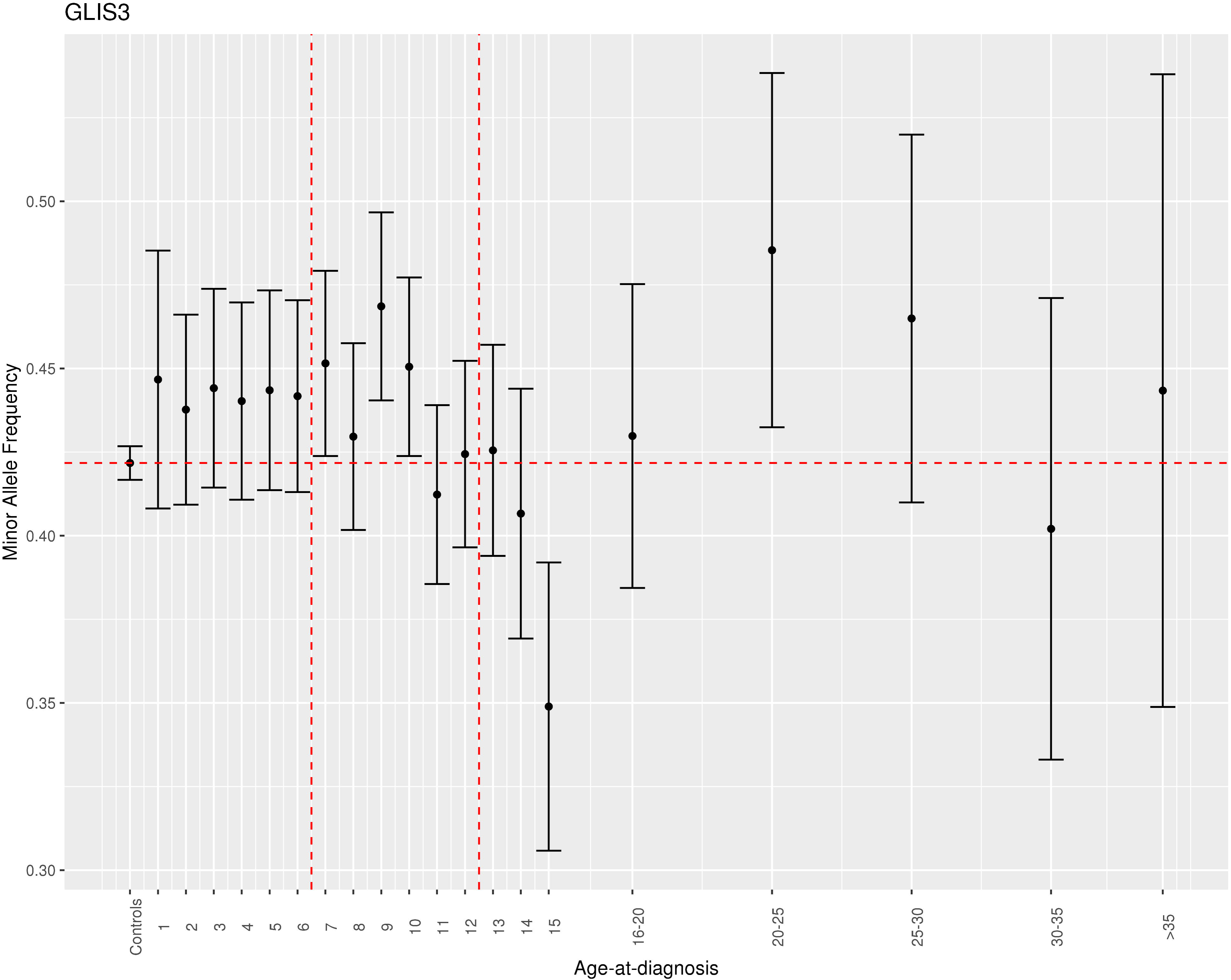
Minor allele frequency of the index variant near the *GLIS3* gene for controls and individuals diagnosed at various ages.

**Supplementary Figure 8:**
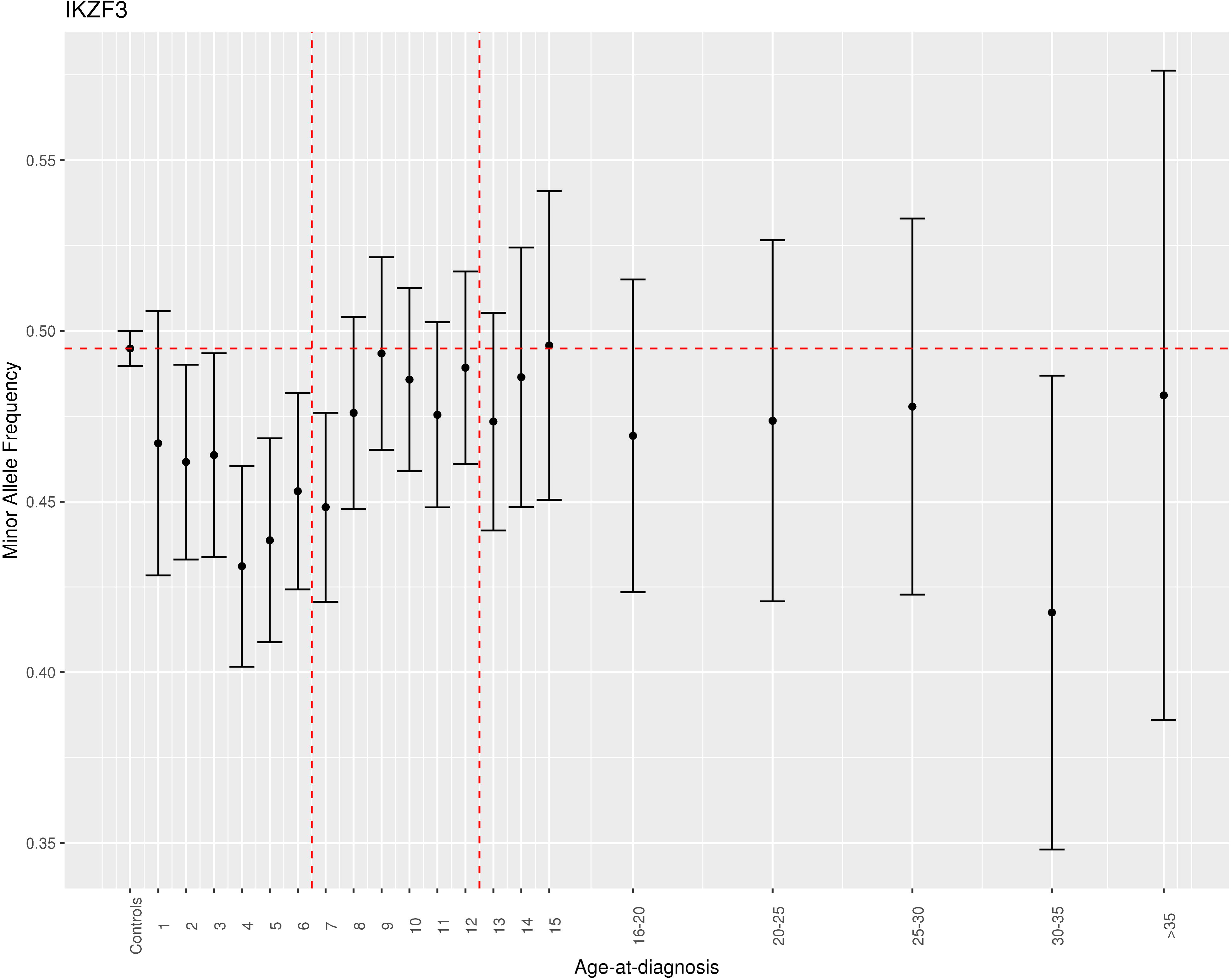
Minor allele frequency of the index variant near the *IKZF3* gene for controls and individuals diagnosed at various ages.

**Supplementary Figure 9:**
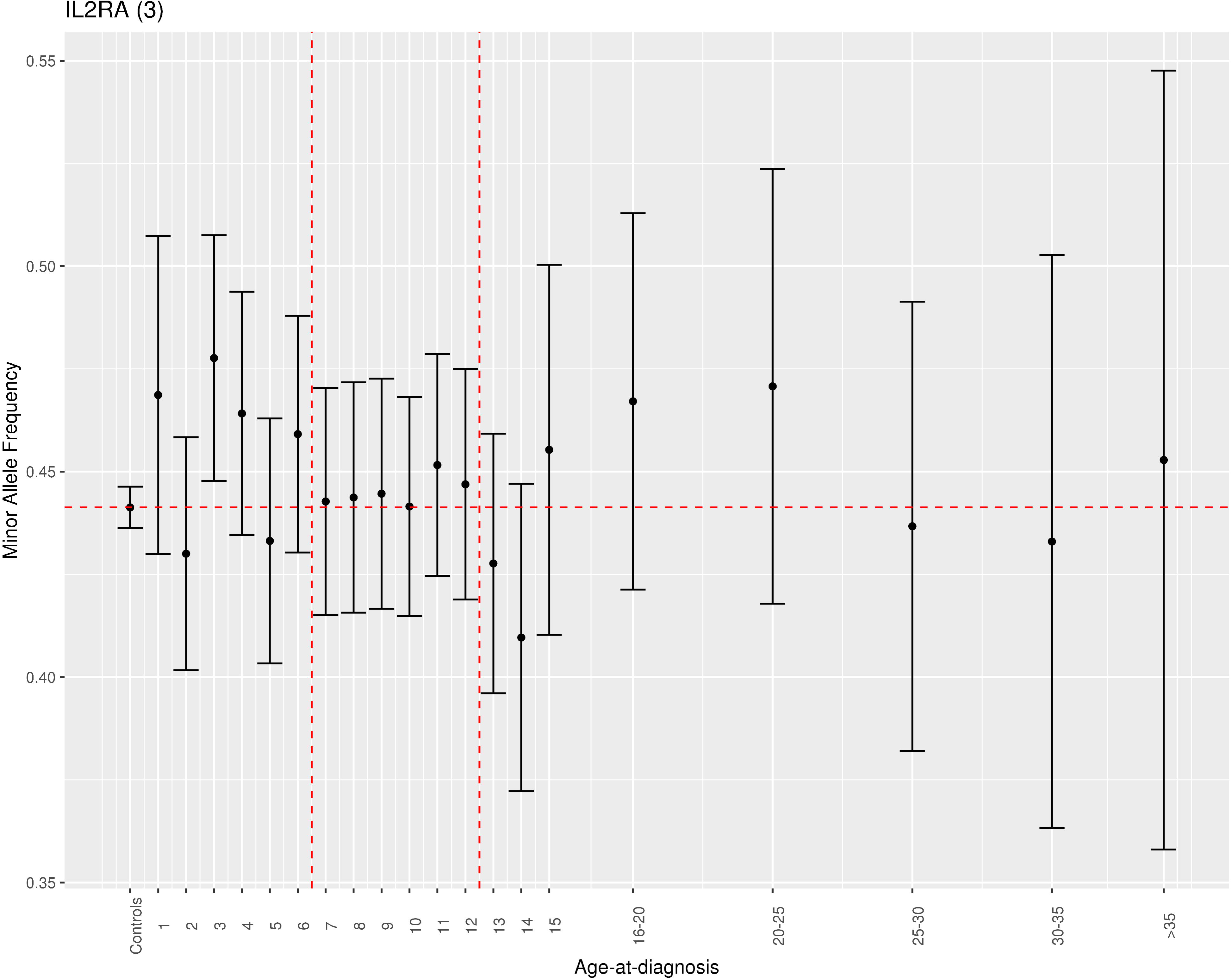
Minor allele frequency of the index variant near the *IL2RA* gene (third index variant) for controls and individuals diagnosed at various ages.

**Supplementary Figure 10:**
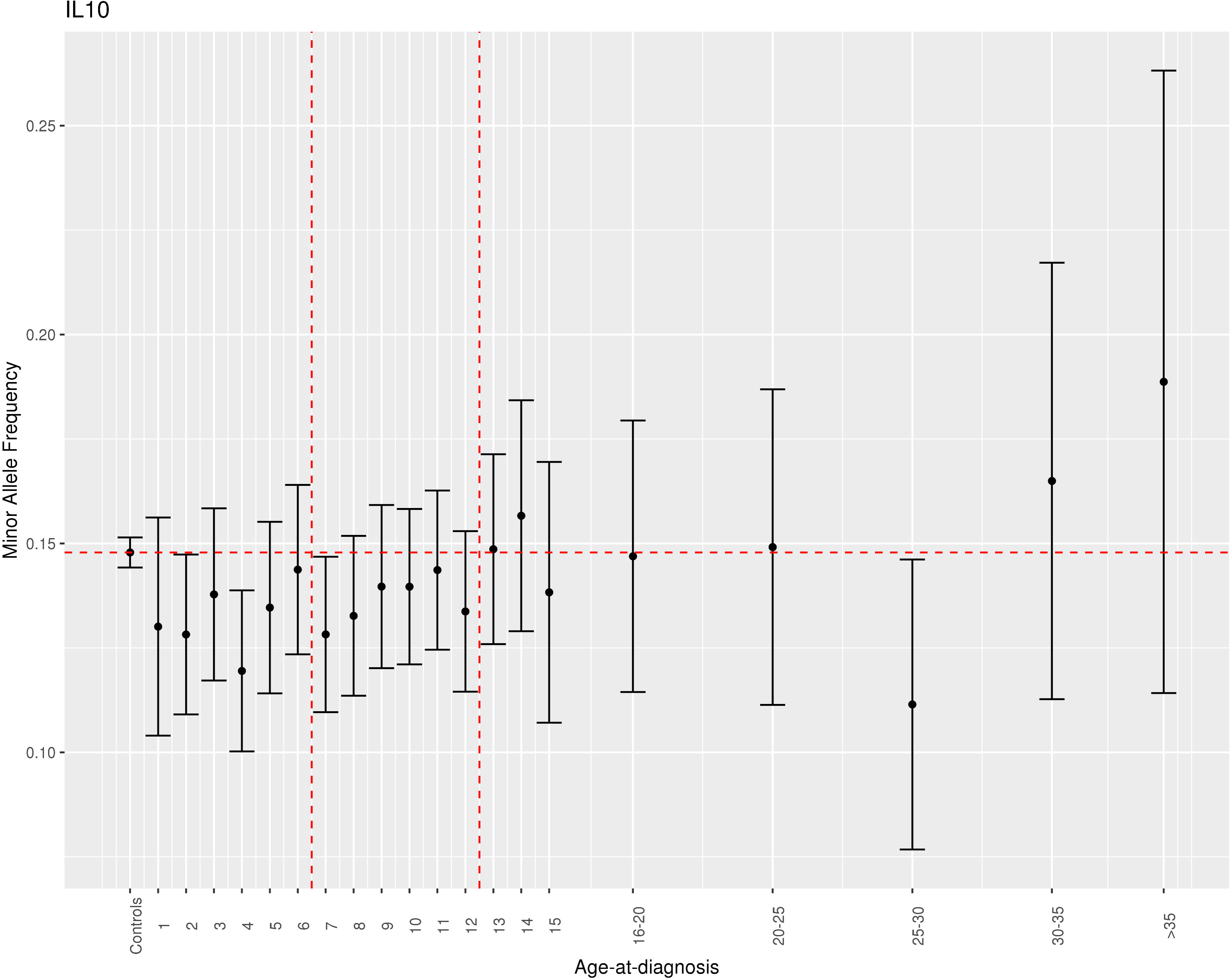
Minor allele frequency of the index variant near the *IL10* gene for controls and individuals diagnosed at various ages.

**Supplementary Figure 11:**
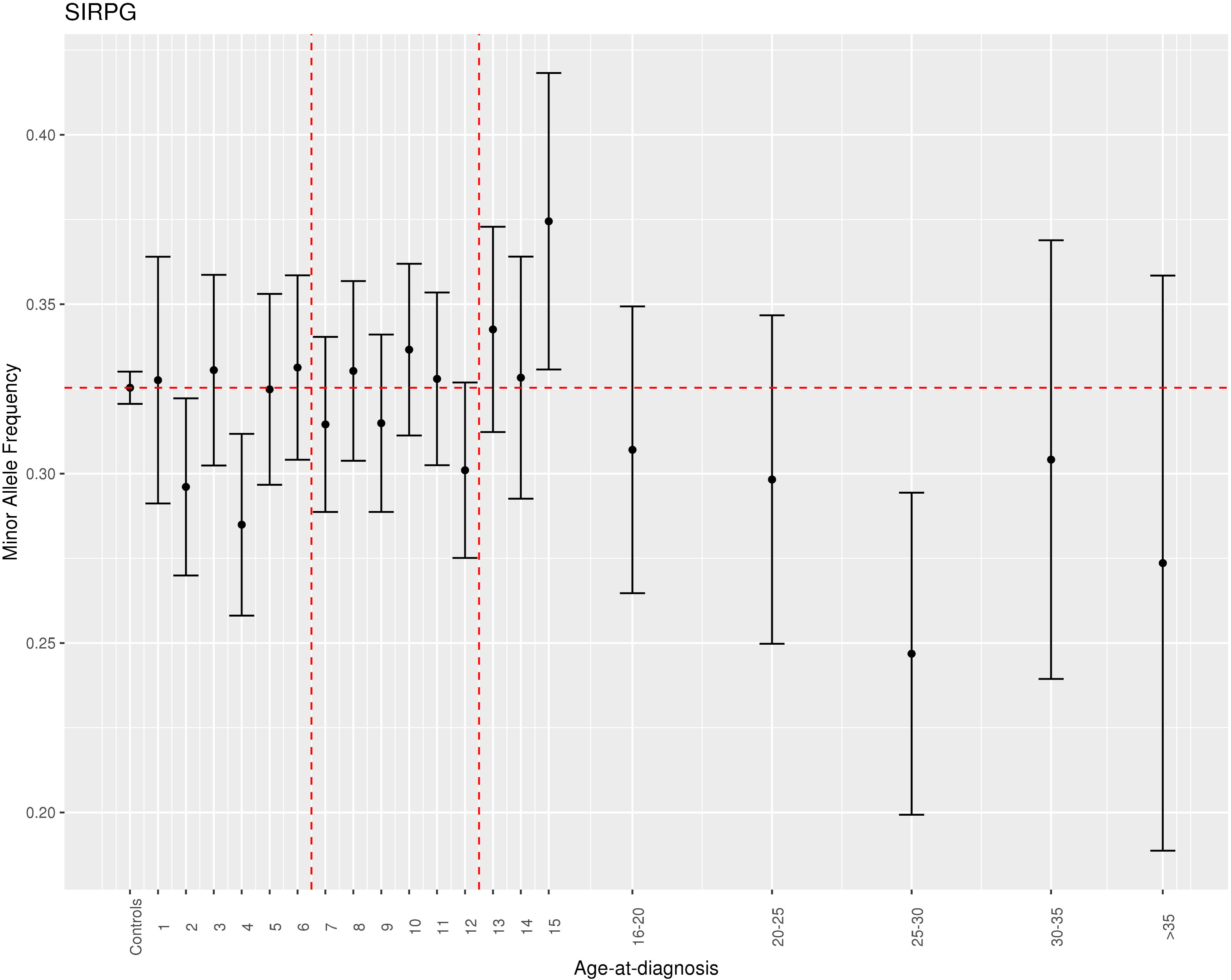
Minor allele frequency of the index variant near the *SIRPG* gene for controls and individuals diagnosed at various ages.

**Supplementary Figure 12:**
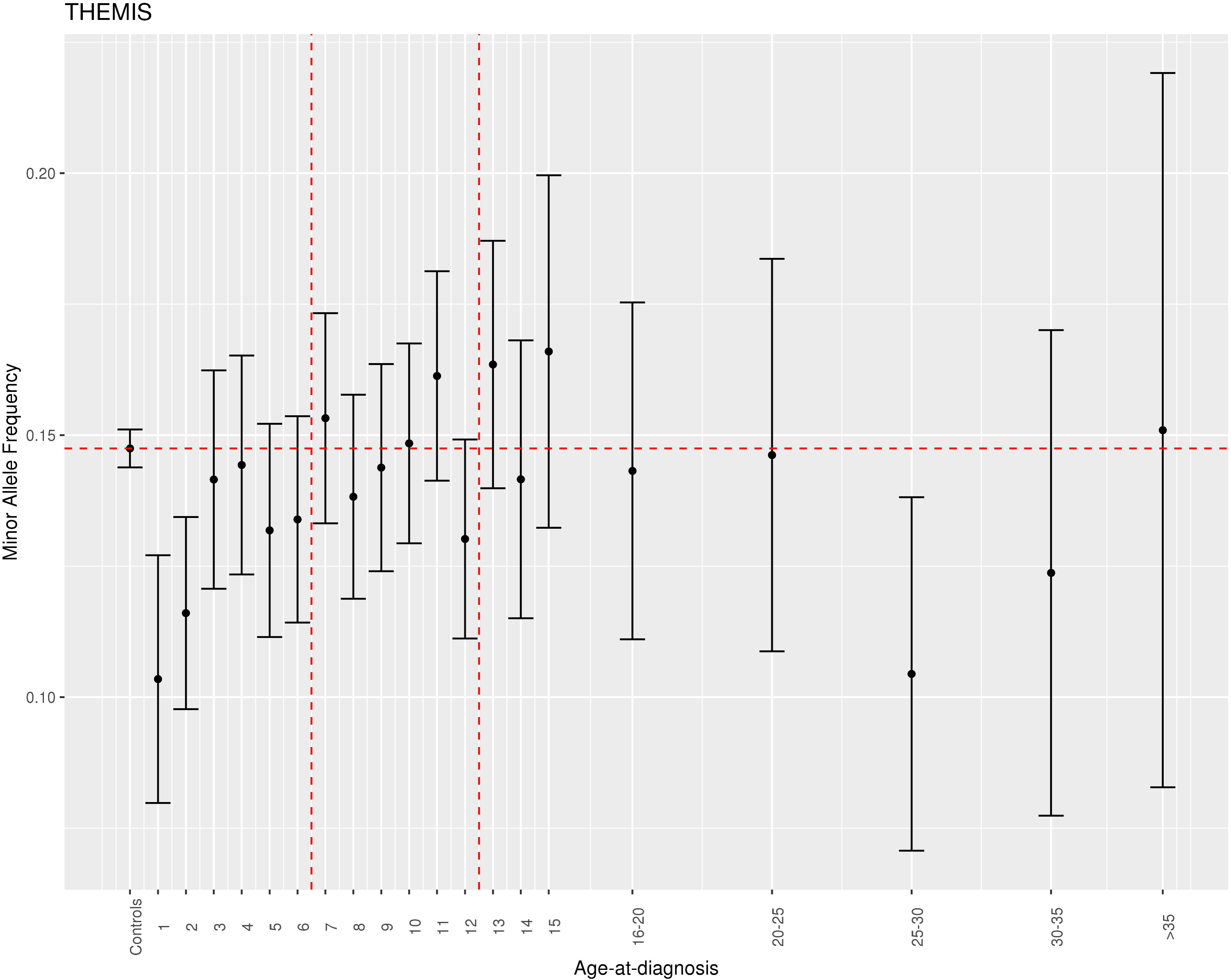
Minor allele frequency of the index variant near the *THEMIS* genes for controls and individuals diagnosed at various ages.

**Supplementary Figure 13:**
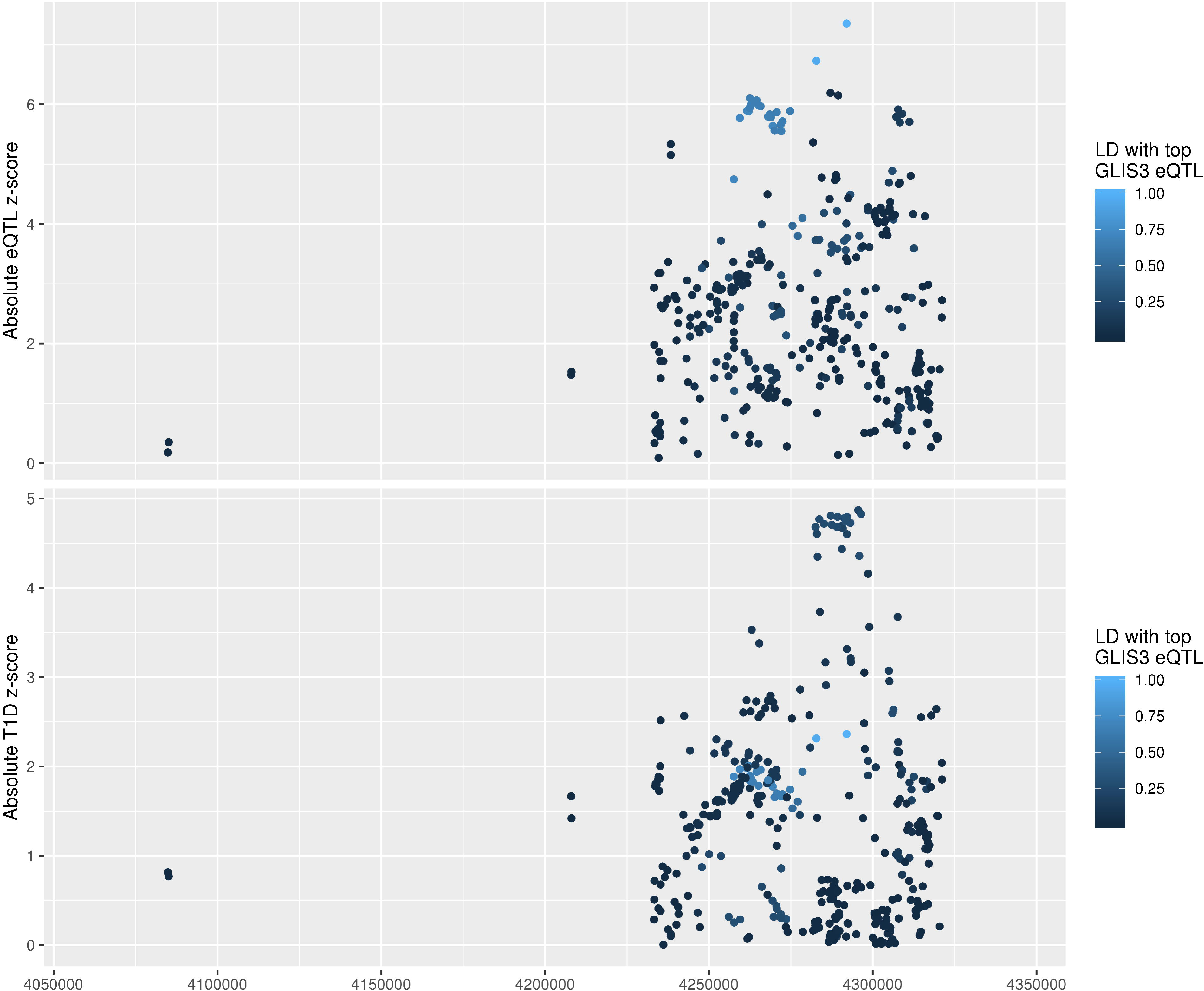
Top panel: association absolute z scores from whole blood eQTL study examining variant effects on GLIS3 mRNA levels, coloured by LD r^2^ to the most strongly associated variant with GLIS3 mRNA expression. Bottom panel: association absolute z scores from logistic regression examining variant associations with type 1 diabetes risk diagnosed at <7 years, using individuals from the UK and Northern Ireland only and adjusting for the five largest principal components as derived from genotype data, coloured by LD r^2^ to the most strongly associated variant with GLIS3 mRNA. Shows most associated disease risk variants are not in high LD with the most associated GLIS3 whole blood eQTLs.

**Supplementary Figure 14:**
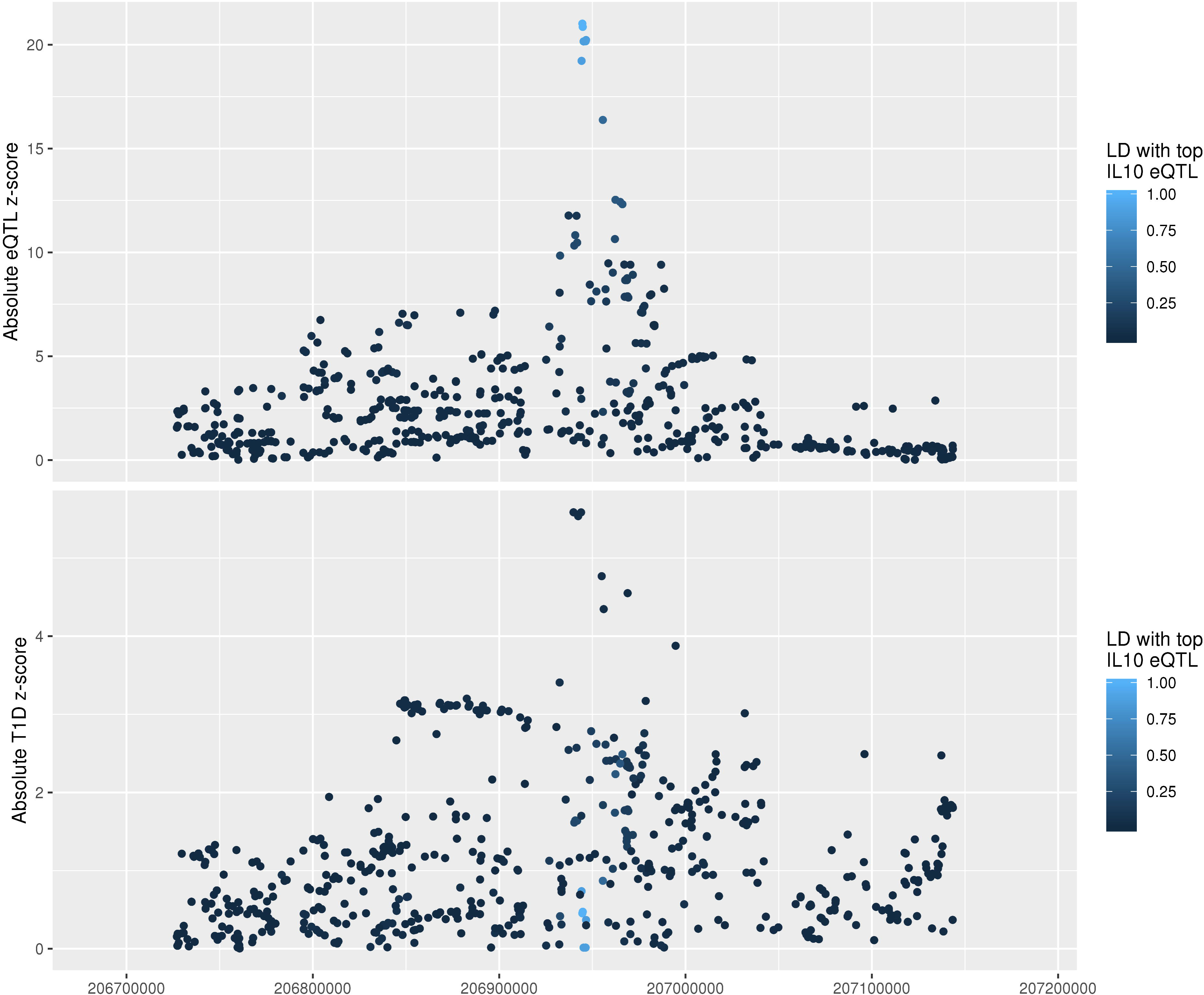
Top panel: association absolute z scores from whole blood eQTL study examining variant effects on IL10 mRNA levels, coloured by LD r^2^ to the most strongly associated variant with IL10 mRNA expression. Bottom panel: association absolute z scores from logistic regression examining variant associations with type 1 diabetes risk diagnosed at <7 years, using individuals from the UK and Northern Ireland only and adjusting for the five largest principal components as derived from genotype data, coloured by LD r^2^ to the most strongly associated variant with IL10 mRNA. Shows most associated disease risk variants are not in high LD with the most associated IL10 whole blood eQTLs.

**Supplementary Figure 15:**
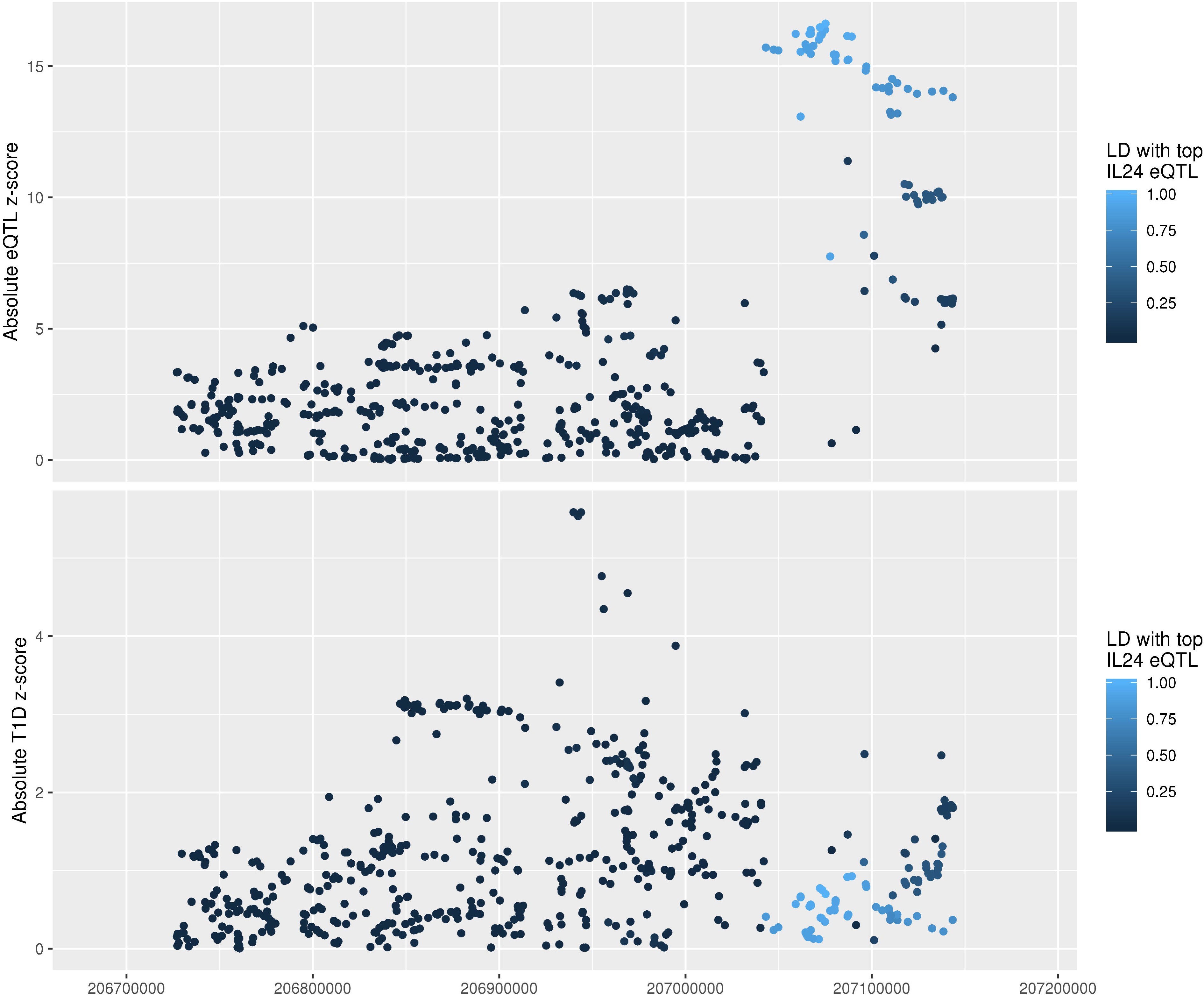
Top panel: association absolute z scores from whole blood eQTL study examining variant effects on IL24 mRNA levels, coloured by LD r^2^ to the most strongly associated variant with IL24 mRNA expression. Bottom panel: association absolute z scores from logistic regression examining variant associations with type 1 diabetes risk diagnosed at <7 years, using individuals from the UK and Northern Ireland only and adjusting for the five largest principal components as derived from genotype data, coloured by LD r^2^ to the most strongly associated variant with IL24 mRNA. Shows most associated disease risk variants are not in high LD with the most associated IL24 whole blood eQTLs.

**Supplementary Figure 16:**
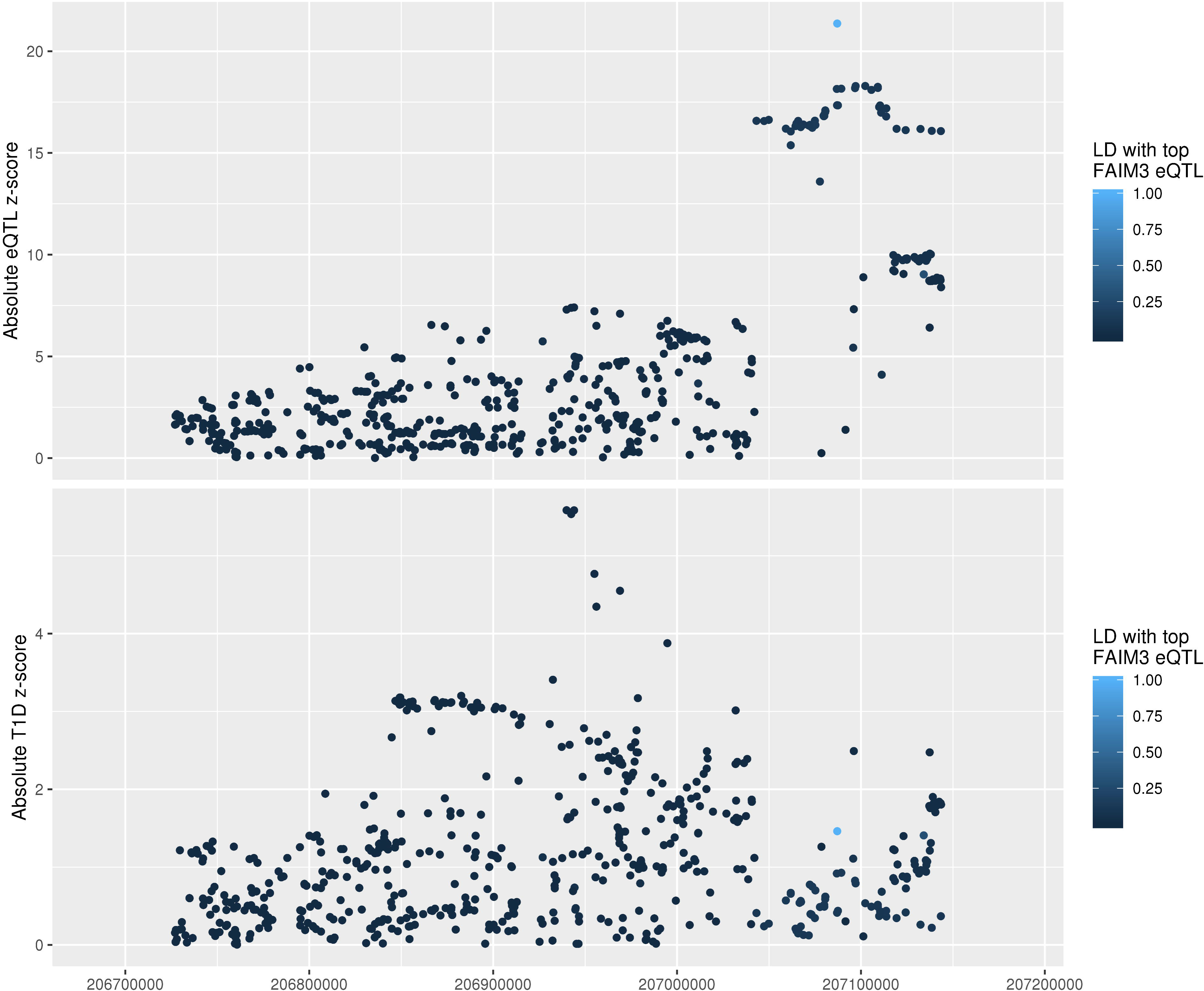
Top panel: association absolute z scores from whole blood eQTL study examining variant effects on FAIM3 mRNA levels, coloured by LD r^2^ to the most strongly associated variant with FAIM3 mRNA expression. Bottom panel: association absolute z scores from logistic regression examining variant associations with type 1 diabetes risk diagnosed at <7 years, using individuals from the UK and Northern Ireland only and adjusting for the five largest principal components as derived from genotype data, coloured by LD r^2^ to the most strongly associated variant with FAIM3 mRNA. Shows most associated disease risk variants are not in high LD with the most associated FAIM3 whole blood eQTLs.

**Supplementary Table 1:** Classical HLA alleles/haplotypes examined in analysis.

**Supplementary Table 2:** Non-HLA variants examined in analysis.

**Supplementary Table 3:** Non-HLA region variants with evidence of heterogeneity in effect size between the <7 and ≥13 groups: Promoter Capture Hi-C (PCHi-C) candidate genes.

**Supplementary Table 4:** Details of non-HLA variants with evidence of heterogeneity in effect size between the <7 and ≥13 groups.

**Supplementary Table 5:** Most likely variants causally associated with type 1 diabetes at the *CTSH* locus from GUESSFM fine mapping analysis.

**Supplementary Table 6:** Most likely variants causally associated with type 1 diabetes at the *GLIS3* locus from GUESSFM fine mapping analysis.

**Supplementary Table 7:** Most likely variants causally associated with type 1 diabetes at the *IKZF3* locus from GUESSFM fine mapping analysis.

**Supplementary Table 8:** Most likely variants causally associated with type 1 diabetes at the *IL2RA* locus from GUESSFM fine mapping analysis. Showing only groups with group posterior probability of greater than 0.9.

**Supplementary Table 9:** Most likely variants causally associated with type 1 diabetes at the *IL10* locus from GUESSFM fine mapping analysis.

**Supplementary Table 10:** Most likely variants causally associated with type 1 diabetes at the *SIRPG* locus from GUESSFM fine mapping analysis.

**Supplementary Table 11:** Most likely variants causally associated with type 1 diabetes at the *THEMIS* locus from GUESSFM fine mapping analysis.

**Supplementary Table 12**: Chip heritability estimates by age-at-diagnosis group under various disease prevalence assumptions.

