## Supplementary Material for "Specific type 1 diabetes risk genes underpin age-at-diagnosis and indicate joint defects in immunity, beta-cell fragility and responses to viral infections in early-onset disease"

**Supplementary Methods**

**Study populations**

The individuals were recruited into various collections, UK-based individuals from the type 1 diabetes genetics consortium (T1DGC) ^1^, the genetic resource investigating diabetes (GRID), the Warren cohort ^2^, the northern Irish GRID ^3^, the 1958 birth control cohort and the UK blood service ^1^, individuals from central Europe, Asia-Pacific and the USA from the T1DGC and individuals from Finland from the IDDMGEN and T1DGEN cohorts ^4^.

**Genotyping and quality control**

All samples were genotyped using the Illumina ImmunoChip ^1^, a specialised platform which densely genotypes regions of interest for autoimmune diseases. Quality control was performed by excluding variants with a minor allele frequency <0.01, a Hardy-Weinberg equilibrium exact test p value of <5×10^-4^ or greater than 5% missing values. Eleven T1D-associated variants we examined failed such quality control and were thus imputed for analysis.

**Relatedness calculations**

We grouped the individuals into three groups, individuals from the UK, individuals from Finland and individuals recruited into the T1DGC study. From each group, we removed those related to closer than second degree by pruning variants correlated to r^2^ >0.2 and removing the major histocompatility (MHC) region, then calculating relatedness using the KING 2.0.9 software ^5^, leaving the total of 27,075 individuals for analysis. After removing variants in linkage disequilibrium (LD) and the MHC region (chromosome 6, positions 25000000-35000000 genome build 36), 28,226 variants were used for relationship inference in the UK group, 28,308 in the T1DGC group and 26,584 in the Finnish group.

**Principal components analysis**

The top 10 genetic principal components were adjusted for in each analysis and were generated using ImmunoChip data filtered so that a set of independent variants (r^2^<0.2) were kept and the MHC region was excluded entirely (chromosome 6, positions 25000000-35000000 genome build 36), leaving a total of 24,456 variants for which to calculate principal components from. The PLINK software was used to carry out this procedure ^6^. For the UK-specific sensitivity analyses and fine-mapping analyses, the top 5 principal components were used as calculated using UK-ancestry individuals only, since there was very little population structure expected and observed in this population.

**False discovery rate**

The false discovery rate used throughout this manuscript is the Benjamini-Hochberg false discovery rate ^7^.

**Binomial test** **examining variants not associated to FDR <0.1**

Given there are 3 AAD groups in our analysis, there are 6 possible ways the log-odds ratio estimates could be ordered. If there were to be no differences in genetic effect between AAD groups, we would expect approximately one sixth of the variants tested to be in each of the possible orders. We therefore performed a binomial test comparing the number of times the ordering was the largest effect estimate in the <7s, the intermediate effect estimate in the 7-13s and the smallest effect estimate in the ≥13s to that expected by chance (1/6=0.167).

**Fine-mapping**

To guard against identifying false positive associations due to imputation, we removed variants from the stochastic search that had a certainty call rate of <75%, a minor allele frequency of <0.005, an absolute Hardy-Weinberg equilibrium z score of >8 (in controls only) and a variant imputation information score of <0.8. We also excluded variants with a difference in imputation information score between cases and controls of more than 0.01, a concordance rate between genotypes and imputed genotype call of <0.8 or where the ratio of the variance to approximate expected variance for a variant with that minor allele frequency (MAF) (≈2×MAF×(1-MAF)) was greater than 0.05.

We adjusted for the top five principal components calculated from UK individuals genotype data, set the prior mean number of expected causal variants in the region to be three and saved the top 10,000 most visited models for downstream analysis.

**Colocalisation analyses**

Summary statistics from both association studies (disease risk and eQTL) were converted to approximate Bayes Factors (ABFs). Posterior probabilities were then calculated for each of the following scenarios: i) association with disease only, ii) association with eQTL only, iii) association with disease and eQTL but distinct signals or iv) shared disease and eQTL signal in the region ^8^. We focus on scenario iv), colocalisation of disease and eQTL signal, which is calculated by multiplying the ABFs for disease and eQTL together for each variant (to get the likelihood of colocalisation) and multiplying this by the prior probability of colocalisation, which we took to be the product of the prior probability of association of each variant for disease and eQTL, set at 1×10^-3^, meaning the prior probability for colocalisation was 1×10^-6^. Dividing this product by the normalising factor, the overall probability of all combinations of each of the four hypotheses and also the probability of no association with disease or eQTL, gives the posterior probability of colocalisation.

**Supplementary Table 1:** Classical HLA alleles/haplotypes examined in analysis.

| **Haplotype/allele name** | **Haplotype/allele definition** | **Individuals included** | **Conditioned on** | **Enough individuals to include?** |
| --- | --- | --- | --- | --- |
| DR3-DQ2 | DRB1*03:01-DQB1*02:01 | Non-DR3/4 only | A*02:01, A*24:02, B*39:06, B*18:01 | Yes |
| DR4-DQ8 | DRB1*04:01-DQB1*03:02  DRB1*04:02-DQB1*03:02  DRB1*04:04-DQB1*03:02  DRB1*04:05-DQB1*03:02  DRB1*04:08-DQB1*03:02  DRB1*04:01-DQB1*03:04  DRB1*04:02-DQB1*03:04  DRB1*04:04-DQB1*03:04  DRB1*04:05-DQB1*03:04  DRB1*04:08-DQB1*03:04  DRB1*04:01-DQB1*02:02  DRB1*04:02-DQB1*02:02  DRB1*04:04-DQB1*02:02  DRB1*04:05-DQB1*02:02  DRB1*04:08-DQB1*02:02 | Non-DR3/4 only | A*02:01, A*24:02, B*39:06, B*18:01 | Yes |
| DR3-DQ2/DR4-DQ8 | Any combination of rows 1 and 2 above, one on either chromosome. | All individuals | A*02:01, A*24:02, B*39:06, B*18:01 | Yes |
| DRB1*13:03-DQB1*03:01 | DRB1*13:03-DQB1*03:01 | Non-DR3/4 only |  | No |
| DRB1*11:04-DQB1*03:01 | DRB1*11:04-DQB1*03:01 | Non-DR3/4 only |  | No |
| DRB1*15:01-DQB1*06:02 | DRB1*15:01-DQB1*06:02 | Non-DR3/4 only |  | Yes |
| DRB1*07:01-DQB1*03:03 | DRB1*07:01-DQB1*03:03 | Non-DR3/4 only |  | Yes |
| DRB1*14:01-DQB1*05:03 | DRB1*14:01-DQB1*05:03 | Non-DR3/4 only |  | No |
| A*02:01 | A*02:01 | All individuals | DR3-DQ2, DR4-DQ8, DR3-DQ2/DR4-DQ8, B*39:06, B*18:01 | Yes |
| A*24:02 | A*24:02 | All individuals | DR3-DQ2, DR4-DQ8, DR3-DQ2/DR4-DQ8, B*39:06, B*18:01 | Yes |
| A*11:01 | A*11:01 | All individuals | DR3-DQ2, DR4-DQ8, DR3-DQ2/DR4-DQ8, B*39:06, B*18:01 | Yes |
| A*32:01 | A*32:01 | All individuals | DR3-DQ2, DR4-DQ8, DR3-DQ2/DR4-DQ8, B*39:06, B*18:01 | Yes |
| DPB1*03:01 | DPB1*03:01 | All individuals | DR3-DQ2, DR4-DQ8, DR3-DQ2/DR4-DQ8, A*02:01, A24*02, B*39:06, B*18:01 | Yes |
| DPB1*04:02 | DPB1*04:02 | All individuals | DR3-DQ2, DR4-DQ8, DR3-DQ2/DR4-DQ8, A*02:01, A24*02, B*39:06, B*18:01 | Yes |
| B*18:01 | B*18:01 | All individuals | DR3-DQ2, DR4-DQ8, DR3-DQ2/DR4-DQ8, A*02:01, A24*02, B*18:01 | Yes |
| B*39:06 | B*39:06 | All individuals | DR3-DQ2, DR4-DQ8, DR3-DQ2/DR4-DQ8, A*02:01, A24*02, B*18:01 | Yes |
| B*44:03 | B*44:03 | All individuals | DR3-DQ2, DR4-DQ8, DR3-DQ2/DR4-DQ8, A*02:01, A24*02, B*18:01 | Yes |

**Supplementary Table 2:** Non-HLA variants examined in analysis.

| **Variant name** | **Chr** | **Position (genome build 37)** | **Publication(s) where association identified*** | **Locus name in Figures in this manuscript** |
| --- | --- | --- | --- | --- |
| rs2476601 | 1 | 114377568 | Onengut-Gumuscu | *PTPN22* |
| rs78037977 | 1 | 172681031 | Fortune | *TNFSF4*** |
| rs6691977 | 1 | 200814959 | Onengut-Gumuscu | *CAMSAP2* |
| rs3024505 | 1 | 206939904 | Onengut-Gumuscu | *IL10* |
| rs4849135 | 1 | 111615079 | Onengut-Gumuscu | *ACOXL* |
| rs13415583 | 2 | 100764087 | Onengut-Gumuscu | *AFF3* |
| rs2111485 | 2 | 163110536 | Onengut-Gumuscu | *IFIH1 (1)* |
| rs35667974 | 2 | 163124637 | Onengut-Gumuscu | *IFIH1 (2)* |
| rs72871627 | 2 | 163136942 | Onengut-Gumuscu | *IFIH1 (3)* |
| rs3087243 | 2 | 204738919 | Onengut-Gumuscu | *CTLA4* |
| rs113010081 | 3 | 46457412 | Onengut-Gumuscu | *CCR5* |
| rs6819058 | 4 | 123114622 | Burren | *IL2/IL21 (1)* |
| rs67797421 | 4 | 123116177 | Burren | *IL2/IL21 (2)* |
| rs2611215 | 4 | 166574267 | Onengut-Gumuscu | *CPE* |
| rs11954020 | 5 | 35883251 | Onengut-Gumuscu | *IL7R* |
| rs72975913 | 6 | 128293932 | Inshaw | *THEMIS* |
| rs72928038 | 6 | 90976768 | Onengut-Gumuscu | *BACH2* |
| rs1538171 | 6 | 126752884 | Onengut-Gumuscu | *CENPW* |
| rs62447205 | 7 | 50465830 | Onengut-Gumuscu | *IKZF1* |
| rs10277986 | 7 | 51028987 | Onengut-Gumuscu | *COBL* |
| rs6476839 | 9 | 4290823 | Onengut-Gumuscu | *GLIS3* |
| rs61839660 | 10 | 6094697 | Wallace | *IL2RA (1)* |
| rs11594656 | 10 | 6122009 | Wallace | *IL2RA (2)* |
| rs6602437 | 10 | 6130077 | Wallace | *IL2RA (3)* |
| rs41295121 | 10 | 6129643 | Wallace | *IL2RA (4)* |
| rs12416116 | 10 | 90035654 | Onengut-Gumuscu | *PTEN*** |
| rs689 | 11 | 2182224 | Onengut-Gumuscu | *INS (1)* |
| rs72853903 | 11 | 2198665 | Onengut-Gumuscu | *INS (2)* |
| rs917911 | 12 | 9905851 | Onengut-Gumuscu | *CD69* |
| rs705705 | 12 | 56435504 | Onengut-Gumuscu | *IKZF4* |
| rs653178 | 12 | 112007756 | Onengut-Gumuscu | *SH2B3* |
| rs9585056 | 13 | 100081766 | Onengut-Gumuscu | *GPR183* |
| rs1456988 | 14 | 98488007 | Onengut-Gumuscu | *LINC01550* |
| rs56994090 | 14 | 101306447 | Onengut-Gumuscu | *MEG3* |
| rs72727394 | 15 | 38847022 | Onengut-Gumuscu | *RASGRP1* |
| rs34593439 | 15 | 79234957 | Onengut-Gumuscu | *CTSH* |
| rs151234 | 16 | 28505660 | Onengut-Gumuscu | *IL27* |
| rs12927355 | 16 | 11194771 | Onengut-Gumuscu | *DEXI (1)* |
| rs193778 | 16 | 11351211 | Onengut-Gumuscu | *DEXI (2)* |
| rs8056814 | 16 | 75252327 | Onengut-Gumuscu | *CTRB1*** |
| rs12453507 | 17 | 38053207 | Onengut-Gumuscu | *IKZF3* |
| rs757411 | 17 | 38775150 | Onengut-Gumuscu | *CCR7* |
| rs1052553 | 17 | 44073889 | Onengut-Gumuscu | *MAPT* |
| rs1893217 | 18 | 12809340 | Onengut-Gumuscu | *PTPN2 (1)* |
| rs12971201 | 18 | 12830538 | Onengut-Gumuscu | *PTPN2 (2)* |
| rs1615504 | 18 | 67526644 | Onengut-Gumuscu | *CD226* |
| rs34536443 | 19 | 10463118 | Onengut-Gumuscu | *TYK2 (1)* |
| rs12720356 | 19 | 10469975 | Onengut-Gumuscu | *TYK2 (2)* |
| rs402072 | 19 | 47219122 | Onengut-Gumuscu | *PRKD2* |
| rs516246 | 19 | 49206172 | Onengut-Gumuscu | *FUT2* |
| rs6043409 | 20 | 1616206 | Onengut-Gumuscu | *SIRPG* |
| rs11203202 | 21 | 43825357 | Onengut-Gumuscu | *UBASH3A* |
| rs6518350 | 21 | 45621817 | Onengut-Gumuscu | *ICOSLG* |
| rs4820830 | 22 | 30531091 | Onengut-Gumuscu | *HORMAD2* |
| rs229533 | 22 | 37587111 | Onengut-Gumuscu | *C1QTNF6* |

*Most recent publication listed. Onengut-Gumuscu ^1^, Fortune ^9^, Burren ^10^, Inshaw ^3^, Wallace ^11^.

**Candidate gene different from previous reports based on publically available gene expression (<https://dice-database.org/>) and promoter-capture Hi-C data (<https://www.chicp.org>).

**Supplementary Table 3:** Non-HLA region variants with evidence of heterogeneity in effect size between the <7 and ≥13 groups: Promoter Capture Hi-C (PCHi-C) candidate genes.

| **Locus name/ karyotype band** | **Tag/index variant** | **Candidate causal genes** | **PCHi-C prioritised protein coding genes*** | **PCHi-C prioritised non-protein coding transcripts** |
| --- | --- | --- | --- | --- |
| *IKZF3* /  17q21 | rs12453507 | *IKZF3*  *ORMDL3*  *GSDMB* | *ORMDL3*  *GSDMB*  *ZPBP2* | *-* |
| *CTSH* /  15q25.1 | rs34593439 | *CTSH* | *CTSH*  *BCL2A1* | *-* |
| *GLIS3* /  9p24.2 | rs6476839 | *GLIS3* | *GLIS3*** | *GLIS3-AS1*** |
| *IL2RA* (3) /  10p15.1 | rs6602437 | *IL2RA* | *RBM17*  *IL2RA*  *GDI2*  *PRKCQ*  *ANKRD16* *FAM208B*  *FBX018* | *RP11-536K7.3*  *PRKCQ-AS1* |
| *IL10 /*  1q32.1 | rs3024505 | *IL10*  *FAIM3* | *IL10*  *FCAMR*  *IL20*  *FAIM3*  *PIGR*  *CD55*  *IL24*  *IL19* | *-* |
| *THEMIS* /  6q22.33 | rs72975913 | *PTPRK*  *THEMIS* | *PTPRK*  *THEMIS* |  |
| *SIRPG /*  20p13 | rs6043409 | *SIRPG*  *SIRPG-AS1* | *-* |  |

*Inshaw et. al ^12^

** Miguel-Escalada et al. ^13^

**Supplementary Table 4:** Details of non-HLA variants with evidence of heterogeneity in effect size between the <7 and ≥13 groups.

| **Locus name/ karyotype band** | **Tag/index variant** | **Candidate causal genes** | **Tissues expressed*** | **BLUEPRINT** ^14^ **/ DICE** database immune cell types expressed***** | **Function** |
| --- | --- | --- | --- | --- | --- |
| *CTSH* /  15q25.1 | rs34593439 | *CTSH* | Ubiquitous | Macrophages, DCs, T and B cells, monocytes | CTSH is a lysosomal protein, has amino- and endopeptidase activity ^15^and has the potential to generate hybrid peptides ^16^. CTSH has been reported to process pro-surfactant protein B ^17^ and pro-granzyme B ^18^ into their active mature forms.  Proteolytic cleavage of TLR3 by CTSH alters the stability, localization and/or the regulation of TLR3 activity ^19, 20^. CTSH-deficient mice have reduced TLR3 expression and type 1 IFN release in response to poly(I:C) stimulation ^21^ and T1D associated SNPs colocalise with a monocyte eQTL signal ^22^. |
| *GLIS3* /  9p24.2 | rs6476839 | *GLIS3* | Pancreas, thyroid gland, kidney, ovary | - | GLIS3 is a transcriptional regulator ^23^, required for insulin expression ^24^, function and transcriptional program of mature eta cells ^25^. A key molecule regulating the response to unfolded protein stress and protection of eta cells from apoptosis ^26,^ ^25^. |
| *IKZF3* /  17q21 | rs12453507 | *IKZF3*  *ORMDL3*  *GSDMB* | *IKZF3*: Small intestine, spleen  *ORMDL3*: Ubiquitous  *GSDMB*: Ubiquitous | *IKZF3*: thymocytes, B, T and NK cells  *ORMDL3*: thymocytes, T, B, and NK cells, eosinophils  *GSDMB*: thymocytes, T, B and NK cells, eosinophils | *IKZF3 (Ailos)*: A transcriptional regulator and a member of the Ikaros gene family ^27^. IKZF3 regulates B cell activation thresholds and differentiation ^28^ and is required for generation of high affinity plasma cells ^29^.  In T cells, IKZF3 acts as a transcriptional repressor to promote Th17 differentiation ^30^ and can modulate the activity of FoxP3 ^31^.  *ORMDL3*: regulates sphingolipid biosynthesis and expression is increased by inflammatory stimuli ^32^. ORMDL3 is a negative regulator of store operated calcium entry ^33^ and modulates lymphocyte activation and cytokine production ^34^.  *GSDMB*: GSDMB can be cleaved by apoptotic caspases ^35^ and the active form of GSDMB induces pyroptotic cell death in epithelial cells ^36^. |
| *IL2RA* (3) /  10p15.1 | rs6602437 | *IL2RA* | Bone marrow, Lymph nodes, spleen | Thymocytes, T, B, NK | *IL2RA* encodes CD25 the alpha chain of the trimeric high affinity IL-2 receptor. IL-2 signaling controls T cell growth and differentiation, NK cell and activated B cell proliferation and is essential for the survival and function of regulatory T cells (Tregs) ^37,^ ^38^ |
| *IL10 /*  1q32.1 | rs3024505 | *IL10*  *FAIM3* | *IL10:* Spleen  *FAIM3:* Spleen, small intestine | *IL10*: Monocytes, activated T, Treg, macrophages and B cells.  *FAIM3*: T and B cells | *IL10*: Interleukin 10 is a key cytokine that modulates immune and non-immune cell function ^39^. IL10 can be produced by specialized regulatory T ^40^ and B cell populations ^41^.  Primary mechanisms include downregulation of MHC class II and cytokine responses of antigen-presenting cells ^42^ and induction of anergy in T cells ^43^. IL10 acts as a potent growth and differentiation factor for B cells ^44^.  *FAIM3:* Encodes FCmR a receptor for natural IgM ^45^ and involved in B cell homeostasis ^46^, control of IL10 production ^47^ and supports early B cell activation events and plasma cell development ^48^. |
| *THEMIS* /  6q22.33 | rs72975913 | *PTPRK* *THEMIS* | *PTPRK*:  Ubiquitous  *THEMIS*:  Spleen, small intestine, thymus | *PTPRK*: Thymocytes, naïve T cells, B cells  *THEMIS*: Thymocytes, T cells | *PTPRK*: Protein tyrosine phosphatase receptor type kappa may be involved in thymic selection of single positive CD4 T cells ^49^.  *THEMIS*: Thymocyte-expressed molecule involved in selection regulates phosphatase activity (Shp1) in developing thymocytes to enable positive selection ^50,^ ^51^. In peripheral T cells, dependent upon intracellular interacting partners, Themis alters T cell signalling to positively ^52^ or negatively ^53^ control T cell activation through the TCR. |
| *SIRPG /*  20p13 | rs6043409 | *SIRPG*  *SIRPG-AS1* | *SIRPG*: Spleen, small intestine, testis | *SIRPG*: T cell  *SIRPG-AS1:* T cell | *SIRPG*: Signaling receptor protein gamma binds to CD47^54^, a ubiquitously expressed integrin associated protein. Interaction of SIRPG with CD47 controls transendothelial migration of T cells ^55^. The T1D associated variant rs2281808 reduces expression of SIRPG in T cells and modulates the activation threshold and differentiation state of CD8 T cells ^56^. |

*Human protein atlas: https://www.proteinatlas.org

** DICE database: https://dice-database.org/

*** T=T cells, B= B cells, NK = Natural Killer cells, DC= Dendritic cells.

***INCLUDE***:

(i) candidate genes nominated in Onengut-Gumuscu et al ^1^ and Inshaw et al ^12^.

(ii) genes prioritised by promoter capture Hi-C (PCHi-C) in immune cells ^12,^ ^57^, or gene promoter interactions with credible SNPs in islet PCHi-C ^13^.

***FILTER ON***:

(i) expression in immune cells (<https://dice-database.org/>) or islets ^58^

***and/or***

(ii) eQTL in whole blood ^59^ passing a Bonferroni corrected p value threshold (p < 2.78 x 10^-13^).

**Supplementary Table 5:** Most likely variants causally associated with T1D at the *CTSH* locus from GUESSFM fine mapping analysis.

| **Variant** | **Position (gr37)** | **Log-odds ratio** | **Reference allele** | **Effect allele** | **Variant posterior probability** | **Group posterior probability** |
| --- | --- | --- | --- | --- | --- | --- |
| rs12592898 | 79229199 | -0.2696 | G | A | 1.11e-02 | 1 |
| rs60254670 | 79229959 | -0.2609 | TGGCCAGAATG | T | 4.57e-03 | 1 |
| rs12148472 | 79231478 | -0.2723 | T | C | 1.90e-02 | 1 |
| rs34843303 | 79234470 | -0.3252 | T | C | 3.38e-01 | 1 |
| rs34593439 | 79234957 | -0.3319 | G | A | 3.17e-01 | 1 |
| rs2289702 | 79237293 | -0.3401 | C | T | 3.11e-01 | 1 |

**Supplementary Table 6:** Most likely variants causally associated with T1D at the *GLIS3* locus from GUESSFM fine mapping analysis.

| **Variant** | **Position (gr37)** | **Log-odds ratio** | **Reference allele** | **Effect allele** | **Variant posterior probability** | **Group posterior probability** |
| --- | --- | --- | --- | --- | --- | --- |
| rs4380994 | 4282536 | -0.1395 | A | G | 3.49e-02 | 0.9852 |
| rs3892354 | 4282942 | -0.1375 | T | G | 2.38e-02 | 0.9852 |
| rs1574285 | 4283137 | -0.1306 | G | T | 7.53e-03 | 0.9852 |
| rs10974435 | 4283682 | -0.1421 | T | C | 5.30e-02 | 0.9852 |
| rs34494309 | 4284961 | -0.1399 | A | AT | 3.62e-02 | 0.9852 |
| rs57884925 | 4285119 | -0.1407 | G | C | 4.16e-02 | 0.9852 |
| rs7024686 | 4287211 | -0.1432 | G | C | 6.40e-02 | 0.9852 |
| rs7041847 | 4287466 | -0.1403 | A | G | 3.94e-02 | 0.9852 |
| rs7034200 | 4289050 | -0.1394 | A | C | 3.50e-02 | 0.9852 |
| rs10814914 | 4289196 | -0.1427 | T | C | 6.04e-02 | 0.9852 |
| rs10116772 | 4290541 | -0.1334 | A | C | 1.10e-02 | 0.9852 |
| rs10814915 | 4290544 | -0.1403 | T | C | 3.60e-02 | 0.9852 |
| rs6476839 | 4290823 | -0.1402 | T | A | 3.17e-02 | 0.9852 |
| rs6476842 | 4291268 | -0.1428 | C | T | 5.65e-02 | 0.9852 |
| rs7020673 | 4291747 | -0.1419 | G | C | 4.66e-02 | 0.9852 |
| rs10974438 | 4291928 | -0.1465 | C | A | 5.38e-02 | 0.9852 |
| rs10758593 | 4292083 | -0.1384 | A | G | 2.36e-02 | 0.9852 |
| rs7867224 | 4292152 | -0.1429 | A | G | 5.98e-02 | 0.9852 |
| rs10814916 | 4293150 | -0.1409 | C | A | 4.37e-02 | 0.9852 |
| rs34706136 | 4294707 | -0.1485 | TG | T | 6.33e-02 | 0.9852 |
| rs10758594 | 4295583 | -0.1454 | A | G | 8.61e-02 | 0.9852 |
| rs4339696 | 4295880 | -0.1301 | G | T | 7.78e-03 | 0.9852 |
| rs10814917 | 4296430 | -0.1438 | A | G | 6.96e-02 | 0.9852 |

**Supplementary Table 7:** Most likely variants causally associated with T1D at the *IKZF3* locus from GUESSFM fine mapping analysis.

| **Variant** | **Position (gr37)** | **Log-odds ratio** | **Reference allele** | **Effect allele** | **Variant posterior probability** | **Group posterior probability** |
| --- | --- | --- | --- | --- | --- | --- |
| rs2941522 | 37910368 | -0.1772 | C | T | 2.98e-02 | 0.9807 |
| rs12946510 | 37912377 | -0.1528 | T | C | 7.27e-04 | 0.9807 |
| rs72538185 | 37916390 | -0.1769 | A | AT | 2.61e-02 | 0.9807 |
| rs907091 | 37921742 | -0.1794 | C | T | 4.98e-02 | 0.9807 |
| rs907092 | 37922259 | -0.1544 | A | G | 9.56e-04 | 0.9807 |
| rs2952140 | 37928059 | -0.1794 | C | T | 4.99e-02 | 0.9807 |
| rs2313430 | 37929816 | -0.1811 | T | C | 7.08e-02 | 0.9807 |
| rs10445308 | 37938047 | -0.1523 | T | C | 3.63e-04 | 0.9807 |
| rs12942330 | 37939839 | -0.1515 | T | C | 3.25e-04 | 0.9807 |
| rs11658993 | 37940808 | -0.1534 | T | C | 4.32e-04 | 0.9807 |
| rs2952144 | 37960017 | -0.1776 | C | T | 3.54e-02 | 0.9807 |
| rs4795395 | 37962987 | -0.1506 | A | T | 2.83e-04 | 0.9807 |
| rs9909593 | 37970149 | -0.1517 | G | A | 3.27e-04 | 0.9807 |
| rs71152606 | 37975214 | -0.178 | CTTCTA | C | 3.30e-02 | 0.9807 |
| rs9303277 | 37976469 | -0.1762 | T | C | 2.64e-02 | 0.9807 |
| rs3816470 | 37985801 | -0.1722 | G | A | 1.13e-02 | 0.9807 |
| rs4795397 | 38023745 | -0.1482 | G | A | 3.42e-04 | 0.9807 |
| rs11557466 | 38024626 | -0.1566 | T | C | 1.56e-03 | 0.9807 |
| rs11078925 | 38025208 | -0.1571 | C | T | 1.82e-03 | 0.9807 |
| rs34120102 | 38026035 | -0.1576 | A | G | 1.99e-03 | 0.9807 |
| rs11655198 | 38026169 | -0.1778 | T | C | 3.54e-02 | 0.9807 |
| rs11650661 | 38026286 | -0.1777 | T | A | 3.50e-02 | 0.9807 |
| rs11655292 | 38026361 | -0.1776 | G | C | 3.43e-02 | 0.9807 |
| rs12709365 | 38027400 | -0.1567 | G | A | 1.56e-03 | 0.9807 |
| rs13380815 | 38027583 | -0.156 | G | A | 1.43e-03 | 0.9807 |
| rs11557467 | 38028634 | -0.1794 | T | G | 4.59e-02 | 0.9807 |
| rs12936231 | 38029120 | -0.1773 | G | C | 3.13e-02 | 0.9807 |
| rs11870965 | 38030205 | -0.1561 | A | T | 1.44e-03 | 0.9807 |
| rs9903250 | 38031030 | -0.1779 | A | G | 3.52e-02 | 0.9807 |
| rs9905959 | 38031138 | -0.1558 | G | A | 1.27e-03 | 0.9807 |
| rs11658278 | 38031164 | -0.1779 | C | T | 3.22e-02 | 0.9807 |
| rs10852935 | 38031674 | -0.1578 | T | C | 1.93e-03 | 0.9807 |
| rs10852936 | 38031714 | -0.1567 | T | C | 1.56e-03 | 0.9807 |
| rs9891174 | 38031802 | -0.1562 | A | T | 1.43e-03 | 0.9807 |
| rs59716545 | 38031857 | -0.1574 | G | T | 1.79e-03 | 0.9807 |
| rs36095411 | 38031865 | -0.1668 | G | T | 3.05e-03 | 0.9807 |
| rs12939457 | 38032188 | -0.1556 | C | T | 1.24e-03 | 0.9807 |
| rs367998020 | 38032200 | -0.1556 | A | AGTGCAGT | 1.22e-03 | 0.9807 |
| rs34189114 | 38032460 | -0.1568 | T | C | 1.67e-03 | 0.9807 |
| rs35736272 | 38032680 | -0.1579 | C | T | 1.98e-03 | 0.9807 |
| rs1054609 | 38033277 | -0.1572 | C | A | 1.71e-03 | 0.9807 |
| rs9907088 | 38035116 | -0.1572 | A | G | 1.72e-03 | 0.9807 |
| rs36038753 | 38035370 | -0.1596 | T | G | 2.59e-03 | 0.9807 |
| rs35569035 | 38035624 | -0.1578 | T | C | 1.94e-03 | 0.9807 |
| rs9910826 | 38035648 | -0.1562 | G | A | 1.48e-03 | 0.9807 |
| rs71355426 | 38035766 | -0.1577 | G | GAGA | 2.00e-03 | 0.9807 |
| rs9904624 | 38036586 | -0.1558 | G | A | 1.38e-03 | 0.9807 |
| rs4795398 | 38038179 | -0.1545 | T | C | 1.09e-03 | 0.9807 |
| rs12939565 | 38038389 | -0.1762 | T | A | 2.49e-02 | 0.9807 |
| rs12939566 | 38038390 | -0.1542 | T | A | 1.02e-03 | 0.9807 |
| rs71152620 | 38039561 | -0.1739 | T | TAACA | 1.63e-02 | 0.9807 |
| rs12232497 | 38040119 | -0.1554 | C | T | 1.23e-03 | 0.9807 |
| rs12232498 | 38040363 | -0.1527 | C | T | 7.75e-04 | 0.9807 |
| rs12941333 | 38040534 | -0.1526 | T | C | 7.64e-04 | 0.9807 |
| rs2872507 | 38040763 | -0.1543 | A | G | 1.02e-03 | 0.9807 |
| rs9908132 | 38042777 | -0.1735 | A | T | 1.52e-02 | 0.9807 |
| rs9901146 | 38043343 | -0.1747 | A | G | 1.81e-02 | 0.9807 |
| rs12936409 | 38043649 | -0.1543 | T | C | 1.02e-03 | 0.9807 |
| rs12103884 | 38045725 | -0.1754 | T | C | 2.08e-02 | 0.9807 |
| rs9906951 | 38048244 | -0.1749 | C | T | 2.03e-02 | 0.9807 |
| rs12950209 | 38049102 | -0.1744 | C | T | 1.88e-02 | 0.9807 |
| rs12950743 | 38049233 | -0.175 | C | T | 1.95e-02 | 0.9807 |
| rs7359623 | 38049589 | -0.1613 | T | C | 1.54e-03 | 0.9807 |
| rs68122720 | 38050092 | -0.1527 | A | AAG | 7.76e-04 | 0.9807 |
| rs8067378 | 38051348 | -0.1763 | G | A | 2.47e-02 | 0.9807 |
| rs12453507 | 38053207 | -0.164 | G | C | 2.66e-03 | 0.9807 |
| rs11651596 | 38056116 | -0.1481 | C | T | 3.35e-04 | 0.9807 |
| rs12949100 | 38057189 | -0.1497 | A | G | 4.44e-04 | 0.9807 |
| rs8069176 | 38057197 | -0.149 | A | G | 3.90e-04 | 0.9807 |
| rs4795399 | 38061439 | -0.1505 | C | T | 5.08e-04 | 0.9807 |
| rs2305480 | 38062196 | -0.1505 | A | G | 5.05e-04 | 0.9807 |
| rs2305479 | 38062217 | -0.1713 | T | C | 1.05e-02 | 0.9807 |
| rs35196450 | 38062942 | -0.1494 | AC | A | 4.09e-04 | 0.9807 |
| rs56750287 | 38062944 | -0.1499 | C | A | 4.51e-04 | 0.9807 |
| rs11078926 | 38062976 | -0.1505 | A | G | 5.06e-04 | 0.9807 |
| rs883770 | 38063381 | -0.1705 | T | C | 8.92e-03 | 0.9807 |
| rs62067034 | 38063738 | -0.1714 | T | C | 1.10e-02 | 0.9807 |
| rs36000226 | 38063929 | -0.172 | C | T | 1.24e-02 | 0.9807 |
| rs36084703 | 38063980 | -0.1727 | C | CA | 1.40e-02 | 0.9807 |
| rs11078927 | 38064405 | -0.1503 | T | C | 4.92e-04 | 0.9807 |
| rs11078928 | 38064469 | -0.1499 | C | T | 4.64e-04 | 0.9807 |
| rs2290400 | 38066240 | -0.1735 | C | T | 1.59e-02 | 0.9807 |
| rs1008723 | 38066267 | -0.1713 | T | G | 1.05e-02 | 0.9807 |
| rs869402 | 38068043 | -0.1665 | T | C | 4.02e-03 | 0.9807 |
| rs1011082 | 38068514 | -0.1652 | T | C | 3.14e-03 | 0.9807 |
| rs921650 | 38069076 | -0.1671 | G | A | 4.56e-03 | 0.9807 |
| rs921649 | 38069274 | -0.1671 | C | T | 4.77e-03 | 0.9807 |
| rs6503524 | 38069809 | -0.1678 | C | T | 5.43e-03 | 0.9807 |
| rs7216389 | 38069949 | -0.1664 | C | T | 4.02e-03 | 0.9807 |
| rs7216558 | 38070071 | -0.1686 | C | T | 6.17e-03 | 0.9807 |
| rs143385463 | 38071855 | -0.164 | ATTT | A | 2.84e-03 | 0.9807 |
| rs1031458 | 38072173 | -0.1671 | G | T | 4.68e-03 | 0.9807 |
| rs1031460 | 38072247 | -0.1707 | T | G | 9.60e-03 | 0.9807 |
| rs8065777 | 38072402 | -0.1679 | C | T | 5.43e-03 | 0.9807 |
| rs7219923 | 38074518 | -0.1671 | C | T | 4.72e-03 | 0.9807 |
| rs7224129 | 38075426 | -0.1667 | G | A | 4.44e-03 | 0.9807 |
| rs8074437 | 38076137 | -0.1685 | T | G | 6.03e-03 | 0.9807 |
| rs4065275 | 38080865 | -0.167 | A | G | 4.86e-03 | 0.9807 |
| rs12603332 | 38082807 | -0.1633 | T | C | 2.45e-03 | 0.9807 |

**Supplementary Table 8:** Most likely variants causally associated with T1D at the *IL2RA* locus from GUESSFM fine mapping analysis. Showing only groups with group posterior probability of greater than 0.9.

| **Variant** | **Position (gr37)** | **Log-odds ratio** | **Reference allele** | **Effect allele** | **Variant posterior probability** | **Group posterior probability** | **GUESSFM group from Wallace et. al.*** |
| --- | --- | --- | --- | --- | --- | --- | --- |
| rs12722563 | 6069561 | -0.4304 | G | A | 2.16e-04 | 0.9985 | A |
| rs12722552 | 6071347 | -0.4324 | C | T | 1.59e-04 | 0.9985 | A |
| rs12722522 | 6078553 | -0.4831 | G | A | 2.74e-02 | 0.9985 | A |
| rs12722508 | 6088743 | -0.4858 | A | T | 3.60e-02 | 0.9985 | A |
| rs7909519 | 6089841 | -0.4928 | T | G | 1.09e-01 | 0.9985 | A |
| rs61839660 | 6094697 | -0.5347 | C | T | 6.23e-02 | 0.9985 | A |
| rs12722496 | 6096667 | -0.4995 | A | G | 6.57e-01 | 0.9985 | A |
| rs12722495 | 6097283 | -0.4886 | T | C | 1.03e-01 | 0.9985 | A |
| rs41295049 | 6109676 | -0.485 | G | A | 1.71e-03 | 0.9985 | A |
| rs41295061 | 6114660 | -0.4752 | C | A | 4.75e-04 | 0.9985 |  |
| rs41295065 | 6116975 | -0.4765 | G | A | 5.81e-04 | 0.9985 |  |
| rs35285258 | 6118770 | -0.1345 | C | T | 9.83e-01 | 0.9829 | C |
| rs6602437 | 6130077 | 0.1339 | T | C | 9.95e-01 | 0.9952 | E |

*Groups defined in ^11^ named ‘A’-‘F’, of which ‘A’, ‘C’, ‘E’ and ‘F’ were found to be type 1 diabetes associated. The group ‘F’ variants were removed from the stochastic search in this analysis due to difference in imputation quality between cases and controls of >1%.

**Supplementary Table 9:** Most likely variants causally associated with T1D at the *IL10* locus from GUESSFM fine mapping analysis.

| **Variant** | **Position (gr37)** | **Log-odds ratio** | **Reference allele** | **Effect allele** | **Variant posterior probability** | **Group posterior probability** |
| --- | --- | --- | --- | --- | --- | --- |
| rs3024505 | 206939904 | -0.2467 | G | A | 3.58e-01 | 0.9953 |
| rs3024495 | 206942413 | -0.2464 | C | T | 2.75e-01 | 0.9953 |
| rs3024493 | 206943968 | -0.2466 | C | A | 3.57e-01 | 0.9953 |
| rs3122605 | 206955041 | -0.2149 | A | G | 5.29e-03 | 0.9953 |

**Supplementary Table 10:** Most likely variants causally associated with T1D at the *SIRPG* locus from GUESSFM fine mapping analysis.

| **Variant** | **Position (gr37)** | **Log-odds ratio** | **Reference allele** | **Effect allele** | **Variant posterior probability** | **Group posterior probability** |
| --- | --- | --- | --- | --- | --- | --- |
| rs2281808 | 1610551 | -0.1436 | C | T | 3.29e-02 | 0.7234 |
| rs3746722 | 1610939 | -0.1415 | A | G | 2.40e-02 | 0.7234 |
| rs2147336 | 1612279 | -0.142 | A | G | 2.26e-02 | 0.7234 |
| rs2147337 | 1612282 | -0.1444 | G | T | 3.03e-02 | 0.7234 |
| rs1535883 | 1612819 | -0.1423 | G | A | 2.49e-02 | 0.7234 |
| rs6074927 | 1613956 | -0.1332 | G | A | 1.43e-02 | 0.7234 |
| rs2318044 | 1614288 | -0.1308 | C | T | 1.03e-02 | 0.7234 |
| rs6043405 | 1615544 | -0.1531 | C | T | 9.55e-02 | 0.7234 |
| rs6110697 | 1615661 | -0.1523 | C | T | 8.86e-02 | 0.7234 |
| rs6043406 | 1615673 | 0.1835 | A | T | 1.25e-02 | 0.2007 |
| rs62186949 | 1615883 | 0.1809 | A | G | 1.12e-02 | 0.2007 |
| rs6043409 | 1616206 | -0.1612 | G | A | 3.80e-01 | 0.7234 |
| rs77747676 | 1617514 | 0.1838 | CAG | C | 1.92e-02 | 0.2007 |
| rs7272255 | 1625156 | 0.1861 | T | A | 2.45e-02 | 0.2007 |
| rs2023565 | 1644304 | 0.1819 | C | T | 1.81e-02 | 0.2007 |
| rs2250261 | 1657529 | 0.1741 | T | C | 2.79e-02 | 0.2007 |
| rs202548 | 1662999 | 0.1755 | T | G | 2.98e-02 | 0.2007 |
| rs202535 | 1666989 | 0.1738 | A | C | 2.72e-02 | 0.2007 |
| rs202531 | 1669151 | 0.1758 | T | C | 3.03e-02 | 0.2007 |

**Supplementary Table 11:** Most likely variants causally associated with T1D at the *THEMIS* locus from GUESSFM fine mapping analysis.

| **Variant** | **Position (gr37)** | **Log-odds ratio** | **Reference allele** | **Effect allele** | **Variant posterior probability** | **Group posterior probability** |
| --- | --- | --- | --- | --- | --- | --- |
| rs67707912 | 128217774 | -0.2046 | T | C | 1.59e-02 | 0.9695 |
| rs13204742 | 128245765 | -0.2006 | G | T | 1.19e-02 | 0.9695 |
| rs6939352 | 128266250 | -0.1979 | T | G | 3.84e-02 | 0.9695 |
| rs9491889 | 128270067 | -0.1994 | C | T | 4.36e-02 | 0.9695 |
| rs9491890 | 128270123 | -0.1996 | G | A | 4.52e-02 | 0.9695 |
| rs9491891 | 128277151 | -0.202 | A | G | 5.82e-02 | 0.9695 |
| rs147626184 | 128277275 | -0.2023 | C | CAACTTGAAT | 5.43e-02 | 0.9695 |
| rs118097399 | 128278233 | -0.2036 | C | T | 6.67e-02 | 0.9695 |
| rs9491892 | 128280358 | -0.1955 | T | G | 3.33e-02 | 0.9695 |
| rs9482848 | 128280375 | -0.1946 | A | T | 2.89e-02 | 0.9695 |
| rs9491893 | 128280931 | -0.2042 | G | A | 7.16e-02 | 0.9695 |
| rs113297984 | 128286301 | -0.1967 | G | A | 3.26e-02 | 0.9695 |
| rs72973797 | 128286386 | -0.1993 | G | A | 4.53e-02 | 0.9695 |
| rs72973800 | 128287158 | -0.1983 | C | T | 3.74e-02 | 0.9695 |
| rs761332 | 128287848 | -0.1975 | G | A | 3.71e-02 | 0.9695 |
| rs9482849 | 128288536 | -0.1968 | T | C | 3.45e-02 | 0.9695 |
| rs12111314 | 128289214 | -0.2009 | C | T | 4.66e-02 | 0.9695 |
| rs11753289 | 128291681 | -0.1973 | T | G | 3.64e-02 | 0.9695 |
| rs9482850 | 128293506 | -0.2003 | C | T | 4.67e-02 | 0.9695 |
| rs9482851 | 128293634 | -0.2029 | C | T | 5.56e-02 | 0.9695 |
| rs72975913 | 128293932 | -0.1996 | C | A | 4.32e-02 | 0.9695 |
| rs72975916 | 128294055 | -0.1992 | C | T | 4.18e-02 | 0.9695 |
| rs7738609 | 128295502 | -0.1882 | C | T | 1.30e-02 | 0.9695 |
| rs138300818 | 128297022 | -0.1818 | T | TG | 8.23e-03 | 0.9695 |
| rs3901020 | 128297604 | -0.1866 | C | G | 1.20e-02 | 0.9695 |
| rs4510698 | 128297611 | -0.186 | C | T | 1.12e-02 | 0.9695 |

**Supplementary Table 12:** Chip heritability estimates by age-at-diagnosis group under various disease prevalence assumptions.

| **Disease prevalence (%) (<7,7-13,≥13)** | **<7 including MHC (95% CI)** | **7-13 including MHC (95% CI)** | **≥13 including MHC (95% CI)** | **<7 excluding MHC (95% CI)** | **7-13 excluding MHC (95% CI)** | **≥13 excluding MHC (95% CI)** |
| --- | --- | --- | --- | --- | --- | --- |
| 0.4,0.4,0.4 | 0.366  (0.346, 0.386) | 0.301  (0.284, 0.319) | 0.236  (0.209, 0.263) | 0.209  (0.189, 0.229) | 0.178  (0.16, 0.195) | 0.094  (0.073, 0.114) |
| 0.5,0.3,0.3 | 0.383  (0.362, 0.404) | 0.285  (0.268, 0.301) | 0.223  (0.197, 0.248) | 0.219  (0.198, 0.24) | 0.168  (0.152, 0.184) | 0.088  (0.069, 0.108) |
| 0.5,0.2,0.2 | 0.383  (0.362, 0.404) | 0.264  (0.249, 0.279) | 0.207  (0.183, 0.23) | 0.219  (0.198, 0.24) | 0.156  (0.141, 0.171) | 0.082  (0.064, 0.1) |
